# A Mouse Model of SARS-CoV-2-Driven Acute Maladaptive Responses and Chronic Systemic Diseases

**DOI:** 10.1101/2025.08.13.670105

**Authors:** Devin Kenney, Giulia Unali, Anna E. Tseng, Joseph Léger, Mao Matsuo, Aoife K. O’Connell, Christina McCooney, Samantha Good, Jack Norton, Fabiana Feitosa-Suntheimer, Mariano Carossino, Hans P. Gertje, Alexander Klose, Neal Paragas, Kevin P. Francis, Jennifer E. Snyder-Cappione, Anna Belkina, Jochen Welcker, Kenneth Albrecht, Ronald B. Corley, Christelle Harly, Nicholas A. Crossland, Florian Douam

**Author notes:** Correspondence: Florian Douam,; Devin Kenney,; Nicholas Crossland,. These authors equally contributed to the study (Devin Kenney, Giulia Unali, Anna E. Tseng, Nicholas A. Crossland, Florian Douam).

## Abstract

Our understanding of SARS-CoV-2 acute and post-acute pathogenesis is hindered by the lack of adequate small animal models. We present RAB/6N, a mouse model prone to severe disease after exposure to SARS-CoV-2 clinical isolates, with lethal cases showing no widespread brain infection typical of the widely used K18-hACE2 mouse model. Lung viral replication in RAB/6N mice remains steady for several days before a decline in viral titers. Delayed initiation of infection clearance is marked by increased lung T-cell extravasation and type-2 immune responses, leading to maladaptive lung consolidation. While systemic antiviral cytokine responses only correlate with SARS-CoV-2 brain infection in K18-hACE2 mice, they are concomitant with pulmonary immune dynamics in infected RAB/6N mice. Convalescent RAB/6N mice display systemic inflammation and decreased antibody titers against SARS-CoV-2 spike RBD, persistent viral RNA and prolonged lymphoid infiltration in the lungs. These animals also exhibit signatures of multi-organ dysfunction, cognitive impairment, cardiac inflammation, hyper- immunoglobulin production, and various autoimmune disorders, illuminating the molecular correlates of various pathologies associated with post-acute sequelae of COVID-19 (PASC). RAB/6N mice pave the way for dissecting the molecular drivers underlying SARS-CoV-2-induced acute maladaptive responses and subsequent post-acute systemic diseases. This preclinical platform also opens opportunities for the exploration of therapeutic interventions against systemic PASC and for anticipating the emergence of PASC-associated comorbidities.

**One-sentence summary:** We generated a hACE2-transgenic mouse model that develops maladaptive lung immune responses upon acute SARS-CoV-2 infection, leading to fatal outcomes or post-acute systemic disease syndromes in convalescent animals.

## INTRODUCTION

Severe Acute Respiratory Syndrome Coronavirus 2 (SARS-CoV-2) infection in humans can result in COVID-19, which manifests in various clinical outcomes, from mild disease to acute respiratory distress and death. Severe COVID-19 is defined by a prominent immunopathology(*1, 2*) characterized by excessive lung inflammation(*3, 4*), dysregulated interferon, complement and humoral responses(*5–9*), and dysfunctional tissue repair mechanisms(*10*), all of which contribute to a potentially acute respiratory distress syndrome (ARDS). To pass beyond the descriptive nature of human studies, the need for animal models enabling the mechanistic dissection of SARS-CoV-2 pathogenesis and developing frontline antiviral countermeasures has been pressing.

Non-human primates have proven instrumental in evaluating antivirals against SARS-CoV-2. However, they are cost-prohibitive, do not generally replicate severe human disease, and limited reagents are available to conduct detailed immunological studies(*11*). Similarly, hamsters develop only mild disease upon SARS-CoV-2 infection(*12, 13*) and are also hardly amenable to comprehensive immune investigations due to limited reagents. In contrast, mice represent a cost-effective and versatile platform to study SARS-CoV-2 pathogenesis and evaluate countermeasures. However, they are not naturally susceptible to human clinical isolates(*14*) (despite some variant-specific infections with limited replication fitness(*15*)) due to the limited interaction of the mouse ortholog of human angiotensin-converting enzyme 2 (hACE2) with the SARS-CoV-2 Spike protein(*16*). To overcome this limitation, mouse-adapted (MA) strains of SARS-CoV-2 have been developed to induce pulmonary infection and disease in wild-type mice(*17–20*), resulting in mild to high lethality depending on the set of adaptive mutations and the age of animals utilized. Yet, MA strains do not replicate the genetic and pathogenic diversity of SARS-CoV-2 at both the quasi-species and variant levels and induce an accelerated disease course(*17–19*), which may confound the modeling of human-like host responses. MA strains also do not address the need for a mouse model amenable to other hACE2-dependent βCoV, including zoonotic strains with spillover potential(*21*).

Acknowledging that hACE2 binding is a significant determinant of human productive infection for SARS-CoV-2(*22, 23*), hACE2-transgenic mouse models(*24–30*) have emerged as promising platforms to study SARS-CoV-2 and other bat βCoV with pandemic potential through their enhanced susceptibility to infection and disease by these viruses. The most widely used version of such models, referred to as K18-hACE2/6J(*24–27, 31*), incorporates eight, randomly integrated copies of the hACE2 cDNA on chromosome 2(*31*) under the control of the mouse keratin 18 (K18) promoter into the genome of a C57BL/6<u>J</u> mouse strain. However, while this model develops fatal clinical disease, death is not associated with severe pulmonary disease and/or ARDS-like features as seen in humans. Instead, it is caused by widespread brain infection following the resolution of pulmonary infection and inflammation (*26*). This feature has confounded the ability of these models to recapitulate human-like pathogenesis during SARS-CoV-2 infection. While alternative hACE2-expressing mouse models or infection procedures can prevent widespread viral brain infection, they do not associate with any clinical disease(*32, 33*).

The current hACE2-transgenic mouse models are also limited in their ability to model clinically relevant post-acute sequelae of SARS-CoV-2 infection. While up to 400 million persons previously infected with SARS-CoV-2 have experienced the persistence of disease phenotypes for several months to years, also referred to as post-acute sequelae of COVID-19 (PASC), our understanding of the causes and consequences of this multifaceted disease remains elusive(*34, 35*). Pre-clinical PASC studies have leveraged MA strains(*20, 36–39*), low-dose infection of K18- hACE2/6J mice(*40*) and the hamster model(*41*), though none of these models replicate PASC-associated systemic inflammation and its consequences.

Here, we introduce a hACE2-transgenic mouse model, RAB/6N, which addresses several limitations of current mouse models of acute SARS-CoV-2 infection and PASC. RAB/6N mice are susceptible to several SARS-CoV-2 clinical isolates and develop severe, lethal disease upon infection. Importantly, the incidence of viral brain infection is significantly reduced in fatal cases of RAB/6N mice compared to K18-hACE2/6J mice, and infection is regionally restricted when it occurs. Lung infection resolution in RAB/6N mice is characterized by delayed initiation of viral clearance, CD4+ T-cell-skewed immune responses concomitant to elevated systemic IFNγ signatures, and maladaptive lung consolidation. In contrast, K18-hACE2/6J mice exhibit rapid lung infection resolution and high systemic IL-6 concentration that temporally align with widespread brain infection. RAB/6N mice that survive lethal infection display systemic inflammation associated with impaired antibody titers against SARS-CoV-2 spike RBD, and persistent viral RNA in the lung. Using these animals, we unraveled tissue-specific transcriptomic dysregulations associated or not with persistence of viral RNA, and indicative of phenotypes previously reported, or presumed to be associated, with PASC.

Collectively, our study introduces a platform to dissect the mechanisms defining acute and post-acute SARS-CoV-2 pathogenesis. RAB/6N mice also enable the evaluation of PASC therapies and the prediction of underappreciated PASC-associated comorbidities.

## RESULTS

### Generation of RAB/6N mouse model

The most widely used SARS-CoV-2 mouse model (K18-hACE2/6J) was generated via random genomic integration of a human ACE2 cDNA cassette under the control of the human keratin 18 (K18) promoter into the C57BL/6<u>J</u> genetic background, resulting in mice expressing 8-30 copies of the hACE2 gene(*31*). Other hACE2 transgene integration strategies have resulted in a trade-off between widespread brain infection or absence of significant clinical disease upon SARS-CoV-2 infection (**Supplemental Text 1**). To better control hACE2 expression, we inserted the same K18-h*ACE2* cassette used in the K18-hACE2/6J model into the ROSA26 locus (on chromosome 6) of a C57BL/6<u>N</u> strain (C57BL/6NTac), resulting in the insertion of a single copy of the transgene (**Fig. S1**). We refer to this model as **R**OSA26-K18-h**A**CE2-C57**B**L/**6N** or RAB/6N mice. RAB/6N mice were then maintained by back-cross to C57BL/6NTac wild-type mice. All the work presented in this study uses hemizygous RAB/6N mice. Notably, C57BL/6<u>N</u> and C57BL6<u>/J</u> mice exhibit multiple genetic polymorphisms, particularly in mitochondrial function and glucose metabolism(*42–45*), which have been linked to differences in susceptibility to infection and inflammation (**Supplemental Text 2**).

To determine if targeted insertion altered h*ACE2* expression, we performed RT-qPCR analysis for *hACE2* in the lung, heart, kidney, spleen and brain of RAB/6N mice. We found no significant difference in *hACE2* expression level between K18-hACE2/6J and RAB/6N in any tissue except for spleen, for which there was significantly higher expression in RAB/6N compared to K18-hACE2/6J (**Fig. 1A**). Additionally, h*ACE2* expression in alveolar and bronchiolar epithelium in the lung was confirmed using RNA *in situ* hybridization (**Fig. 1B**).

**Figure 1.**
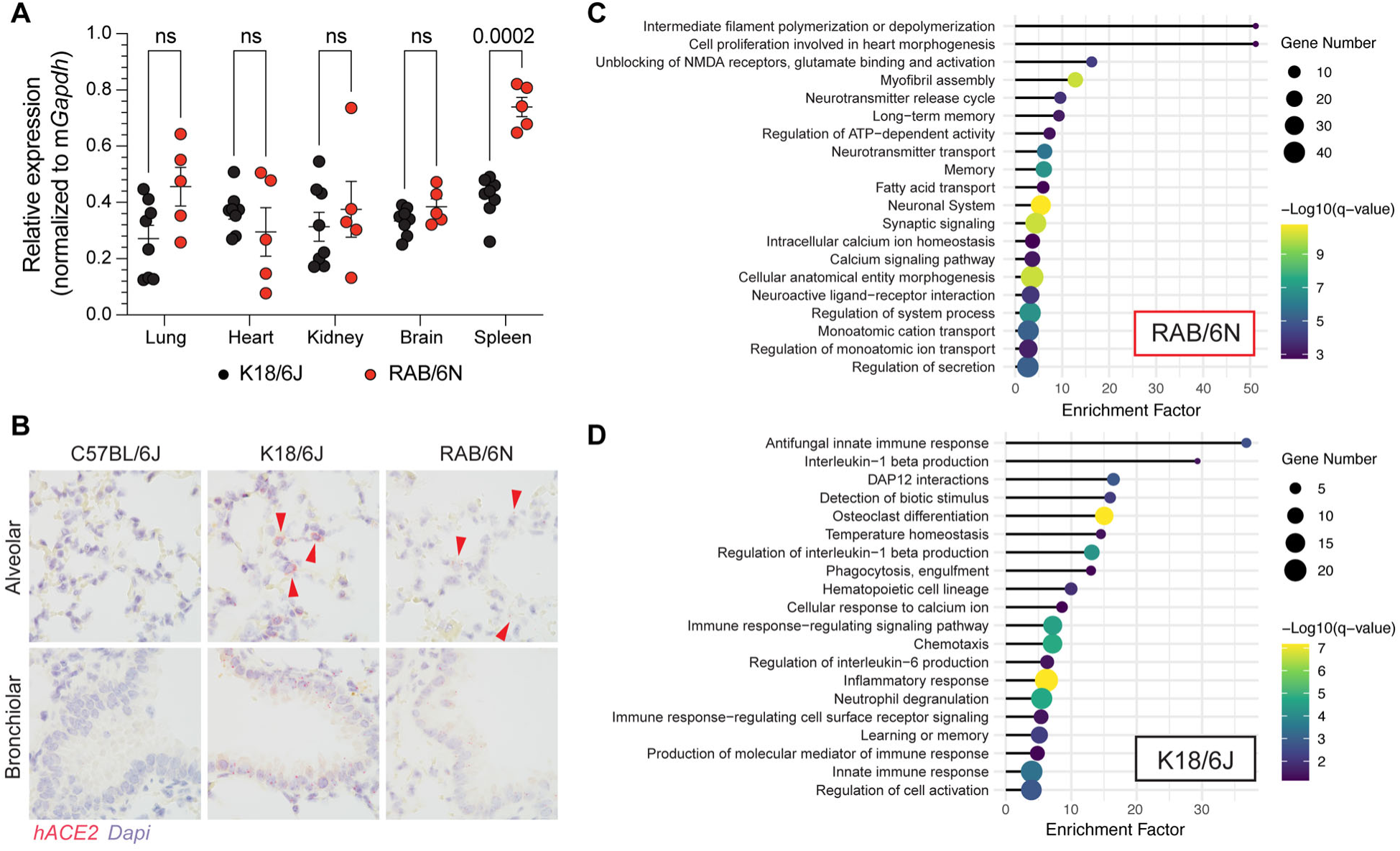
Comparison of RAB/6N and K18-hACE2/6J mice. **(A)** RT-qPCR analysis of *hACE2* expression in lung, heart, kidney, brain and spleen of non-infected RAB/6N (red) and K18- hACE2/6J mice (black). Two-way ANOVA. P-value indicated on graphs. N=5-7. **(B)** RNA fluorescent *in situ* hybridization (FISH) targeting *hACE2* in non-infected C57BL/6J, K18-hACE2/6J and RAB/6N lung tissue. **(C-D)** Pathway enrichment analysis performed using bulk RNA-sequencing in lung tissue from non-infected K18-hACE2/6J **(C)** and RAB/6N **(D)** mice (n=3 male and 4 female).

Bulk RNA-sequencing of total lung homogenates of naive mice revealed 947 genes differentially expressed between RAB/6N and K18-hACE2/6J mice, with 678 genes expressed at higher in RAB6/N mice compared to K18-hACE2/6J (**Fig. S2A)**. Gene pathways enriched in RAB/6N mice were related to neuronal and calcium signaling, molecular transport and cellular morphogenesis (**Fig. 1C**). Genes enriched in K18-hACE2/6J mice associated with immune- related pathways, including innate immune responses, IL1β and IL-6 production, phagocytosis, inflammatory responses and chemotaxis (**Fig. 1D**). Notably, our analysis revealed specific genes displaying both model- and sex-bias expression profiles (**Fig. S2B**). Genes associated with cardiac muscle processes (*Tnnt2, Mybpc3, Hrc, Csrp3, Myh6, Mlip3, and Casq2*), ion-transport channel regulation (*Kcnj11, Kcnj5,* and *Slc17a7*) endothelial to mesenchymal transition (*Nkx2-5, Hand2,* and *Shox2*), and regulation of iron storage in macrophages (*Hamp*) were more enriched in male than female RAB/6N mice (**Fig. S2B**), while genes associated with inflammation (*Il1b, Fgr, Nfam1, Cxcl2, Il23r,* and *Ifi208*), T cell and myeloid cell activation (*Lilrb4a, Csf3r, Sirb1b, Sirb1c,* and *Gm5150*) and neutrophil (*Cd300ld* and *Cd300ld3*) and platelet activation (*Ptafr*) were more enriched in male than female K18-hACE2/6J mice (**Fig. S2B**). Collectively, our analysis suggests that the lungs of RAB/6N and K18-hACE2/6J mice exhibit major transcriptomic differences at steady state.

### RAB/6N mice develop severe disease upon SARS-CoV-2 infection but are less susceptible to fatal outcomes than K18-hACE2/6J mice

To determine susceptibility of RAB/6N mice to SARS-CoV-2 infection, RAB/6N and K18- hACE2/6J mice were intranasally inoculated with varying doses of SARS-CoV-2 ancestral strain (WA-1), Delta (B.1.617.2) and early Omicron (BA.1) variants and assessed for survival and weight loss. Similar to K18-hACE2/6J mice, RAB6/N mice demonstrated dose-dependent disease severity and fatality rate upon infection with the virulent SARS-CoV-2 WA-1 isolate and highly virulent Delta isolate, with no significant difference between males and females (**Fig. 2A, B; Fig. S3A, B**). While a relatively low dose of WA-1 (103 PFU) did not expose major differences in weight loss or survival between the two mouse models, a moderate inoculum (104 PFU) of either WA-1 and Delta showed a reduced susceptibility of RAB/6N to fatal disease compared to K18-hACE2/6J (75% vs. 33% survival; **Fig. 2A, B; Fig. S3A, B**). While increasing the WA-1 inoculum dose (105 PFU) abrogated survival differences between the two models, RAB/6N mice exhibited enhanced survival and recovery from severe disease upon Delta infection at high dose (105 PFU). Notably, upon high dose Delta infection, only a single K18-hACE2/6J mouse survived infection but never developed disease, while RAB/6N mice developed disease and 20% of mice survivedacute disease (**Fig. 2A, B; Fig. S3A, B**). Using our WA-1 survival data, we determined the 50% lethal dose (LD50) of K18-hACE2/6J and RAB/6N mice to be ∼4.7×103 pfu and 1.8×104 pfu, respectively – further highlighting the increased resistance of RAB/6N to fatal outcomes upon acute disease (**Fig. S4**). Notably, while death-associated clinical symptoms in the K18-hACE2/6J model demonstrated evident clinical signs of neurological diseases (i.e., stupor, tremors, proprioceptive defects, and abnormal gait), these symptoms were not apparent in fatal cases of RAB6/N, suggesting a differential susceptibility to brain infection in this model.

**Figure 2.**
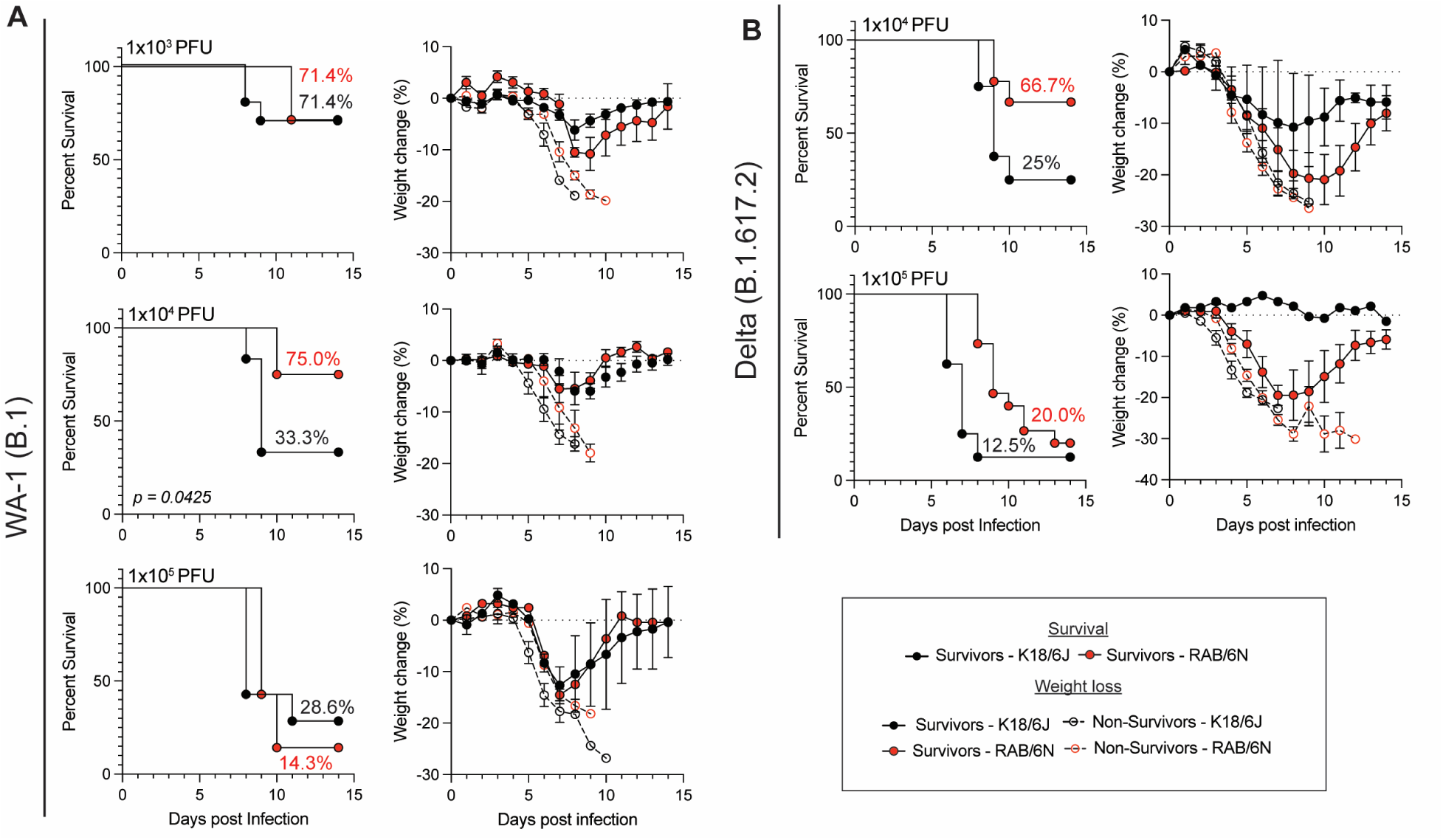
RAB/6N mice develop severe disease upon SARS-CoV-2 infection but are less susceptible to fatal outcomes than K18-hACE2/6J mice. (A-B) 12–22-week-old male and female RAB/6N (red) and K18-hACE2/6J mice (black) were infected intranasally with SARS-CoV- 2 WA-1 **(A)** or Delta **(B)** and monitored for 14 days. **(A)** Mice were infected with 1×103 (top; n=10 K18 and 7 RAB), 1×104 (middle; n=12 K18 and 8 RAB), and 1×105 pfu (bottom; n=7 K18 and 7 RAB) of WA-1 isolate. **(B)** Mice were infected with 1×104 (top; n=8 K18 and 9 RAB) and 1×105 pfu (bottom; n=9 K18 and 14 RAB) of Delta variant. Survival percentage (left) and weight loss (right) are shown. A Mantel-Cox analysis was performed to assess the significance of survival.

Infection with Omicron BA.1 only causes minor disease (mean=3% weight loss at the peak of disease) in K18-hACE2/6J mice when inoculated with a 104 PFU viral dose(*46*), consistent with previous reports. However, strikingly, RAB/6N mice demonstrated increased susceptibility to disease (mean=9% weight loss at the peak of disease) upon Omicron BA.1, although no fatal outcomes were observed for either mouse model (**Fig. S5A)**. Furthermore, weight loss changes in BA.1-infected RAB/6N exhibited a sex-biased profile (**Fig. S5B**), with males displaying more severe disease than females, consistent with human clinical data(*47–49*). These data demonstrate that K18-hACE2/6J and RAB/6N mice elicit differential susceptibility to SARS-CoV- 2 infection in a dose and variant-dependent manner. Notably, attenuated variants (i.e., Omicron BA.1) promote an enhanced disease phenotype in RAB/6N compared to K18-hACE2/6J, suggesting that differential host-pathogen interactions shape clinical disease between these two mouse models.

### Fatal outcomes in RAB6/N mice are uncoupled from widespread brain infection

We then investigated whether, like K18-hACE2/6J, RAB/6N mice are susceptible to widespread SARS-CoV-2 infection of the brain, and whether such an event is associated with fatal outcomes. To do so, we inoculated both mouse strains with 105 PFU of WA-1 isolate, a dose that causes similar lethality in both strains, and collected brain tissues at 2, 4, and 7 dpi. As previously reported(*26*), lethality in K18-hACE2/6J was associated with significant brain infection, with 75% (3:4 mice) and 100% (5:5 mice) of mice displaying brain infection at 4 and 7 dpi, respectively (**Fig. 3A**). In contrast, 60% of RAB/6N (3:5 mice) mice displayed no evidence of infectious viral particles at either time point, despite fatal outcomes. At 7 dpi, K18-hACE2/6J exhibited a statistically significant higher brain viral burden compared to RAB/6N (*p=0.002*), and only one out of five RAB/6N mice displayed a titer greater than 104 PFU/mg tissues (**Fig. 3A**). We also noted a significant increase of infectious virus in the brain between 2 and 4 dpi (*p = 0.0003*) and 4 and 7 dpi (*p = 0.0003)* in K18-hACE2/6J mice, while this was not the case in RAB/6N mice.

**Figure 3.**
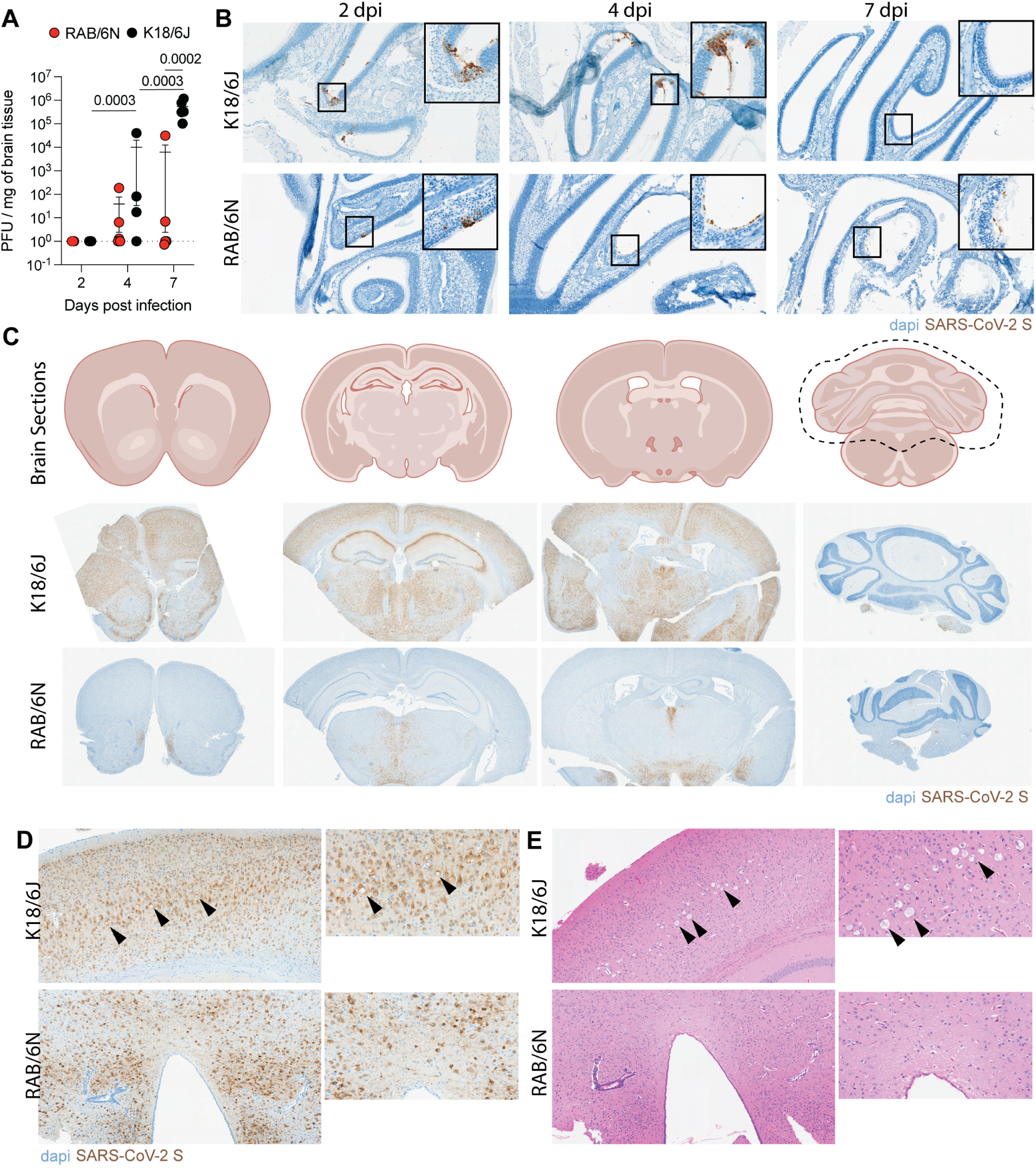
Fatal outcomes in RAB6/N mice are uncoupled from widespread brain infection. **(A)** Viral titers, determined by plaque-forming assay, in brain tissue of K18-hACE2/6J (black) and RAB/6N (red) at 2, 4, and 7-days post-infection. Two-way ANOVA. P-values indicated on graph **(B)** Immunohistochemistry (IHC) for SARS-CoV-2 Spike (S) performed on olfactory epithelium tissue sections from K18-hACE2/6J (top) and RAB/6N (bottom) mice at 2, 4, and 7 dpi. **(C)** IHC for SARS-CoV-2 S performed on coronal brain sections from K18-hACE2/6J (middle panels) and RAB/6N (bottom panels) at 7 dpi. **(D)** IHC for SARS-CoV-2 S and **(E)** hematoxylin and eosin (H&E) staining on brain sections from K18-hACE2/6J (top) and RAB/6N (bottom) at 7 dpi. Neuronal death and brain lesions are represented by black arrows in panel **F**.

The olfactory epithelium (OE) has been suggested as a potential entry route for SARS-CoV-2 neuroinvasion and dissemination(*26*). Immunohistochemistry (IHC) against SARS-CoV-2 spike (S) revealed an increased presence of viral antigen in the OE of K18-hACE2/6J mice at 2 and 4 dpi compared to RAB/6N mice with apparent damage to the epithelial layer (**Fig. 3B**). Notably, while minimal to no viral antigen was observed in K18-hACE2/6J mice at 7 dpi, residual amounts of antigen were detected in RAB/6N mice at 7 dpi (**Fig. 3B**). We also challenged RAB/6N mice with a fatal dose of Delta variant (105 PFU), which has enhanced neuroinvasive properties compared to WA-1 in K18-hACE2/6J mice(*50*), and assessed viral titers in the brain. Ten out of eighteen RAB/6N mice (56%) were free of any infectious virus in the brain at the time of death despite the high viral inoculum (**Fig. S6**), further underscoring that brain infection is not positively correlated with fatal outcome in this model.

Consistent with our viral titer findings, 60% of RAB/6N mouse brains analyzed at 7 dpi following WA-1 infection were negative for S (3:5), while 100% of K18-hACE2 mouse brains were positive (4:4) (**Fig. S6B**). Notably, while S distribution in K18-hACE2/6J brains at 7 dpi was widespread throughout the grey matter of the forebrain (telencephalon and diencephalon), midbrain (mesencephalon) and hindbrain (metencephalon, with the exception of the cerebellum, and myelencephalon), S-positive RAB/6N brain displayed mild and restricted antigen distribution within the thalamus and hypothalamus at 7 dpi (**Fig. 3C**), consistent with some observations in humans (*51, 52*) and hamsters(*53*). Additionally, hematoxylin and eosin (H&E) staining revealed limited histopathology in S-positive regions of the RAB/6N brain at 7 dpi, mainly associated with mononuclear perivascular cuffing and gliosis and, which significantly contrasted with evidence of neuronal degeneration and necrosis (most evident in cortical and hippocampal neurons) in S-positive region of the K18-hACE2/6J brain at that time point, as we previously reported(*26*) (**Fig. 3D, E**). Collectively, our findings highlight that RAB6/N mice display significantly reduced incidence of brain infection compared to K18-hACE2/6J mice. Additionally, when observed, neuroinvasion is regionally restricted in RAB/6N.

### The lungs of RAB/6N mice display distinctive viral replication dynamics and histological outcomes

Lung viral titers at 2 dpi were significantly higher in K18-hACE2/6J infected with 1×105 PFU of WA-1 (2.2×106 ± 7.0×105 PFU/mg of tissue) compared to RAB/6N mice (6.5×103 ± 7.4×103 PFU/mg of tissue) (**Fig. 4A**). K18-hACE2/6J showed a significant reduction in lung viral titers between 2 and 4 dpi (2.2×105 ± 4.8×105 PFU/mg of tissue) and an appreciable, yet non-significant reduction between 4 and 7 dpi (5.6×102 ± 4.4×102 PFU/mg of tissue) that inversely correlates with an increase in brain viral titers (**Fig. 4A, B**). Conversely, RAB/6N mice showed no substantial reduction in lung viral titer between 2 and 4 dpi (4.0×103 ± 2.0×103 PFU/mg of tissue), and an appreciable, but non-statistically significant, reduction between 4 and 7 dpi (11.2 ± 11.6 PFU/mg of tissue) (**Fig. 4A, C**). Infection of RAB/6N mice with a SARS-CoV-2 WA-1 recombinant virus expressing NanoLuc (SARS-CoV-2-NL) in place of ORF7a confirmed the lung-specific tropism of SARS-CoV-2 in RAB/6N mice across all other subperitoneal tissues (**Fig. S7).**

**Figure 4.**
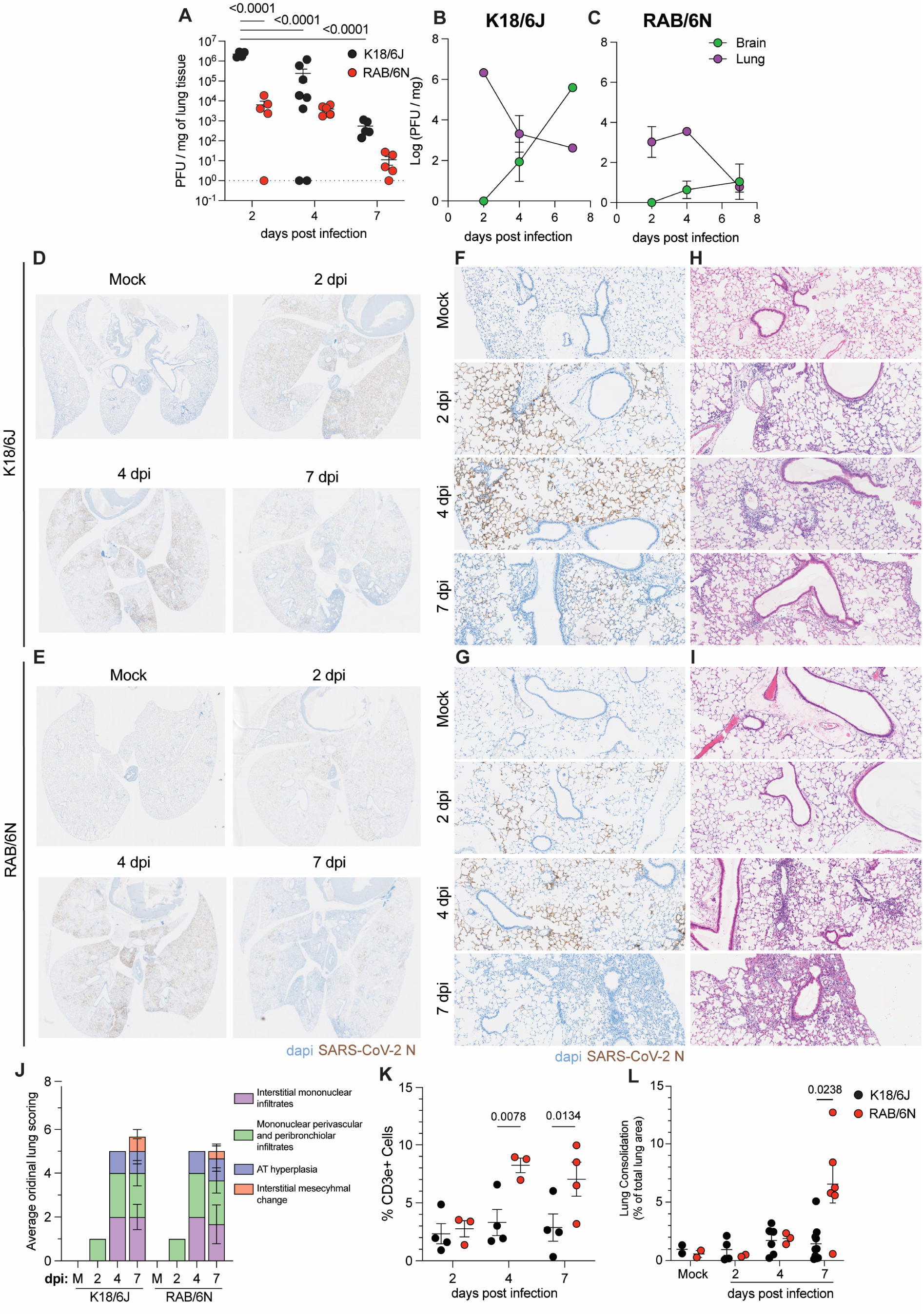
The lungs of RAB/6N mice display specific viral replication dynamics and histopathological features. **(A)** Viral titers, determined by plaque-forming assay, in lung tissue of K18-hACE2/6J (black) and RAB/6N (red) at 2, 4, and 7 days post-infection. Two-way ANOVA. P-values are indicated on the graph. **(B-C)** Line plot displaying mean lung (purple) and brain (green) viral titers at 2, 4, and 7 dpi in **(B)** K18-hACE2/6J and **(C)** RAB/6N mice. **(D-E)** Whole- lung IHC against SARS-CoV-2 nucleocapsid (N) in the lungs of **(D)** K18-hACE2/6J and **(E)** RAB/6N mice at 2, 4, and 7 dpi and in non-infected (mock) mice. **(F, G)** 10x Images showing the distribution of SARS-CoV-2 N antigen in the lung at 2, 4, and 7 dpi in K18-hACE2/6J **(F)** and RAB/6N **(G)** mice. **(H-I)** 10x images of H&E staining on lung sections of **(H)** K18-hACE2/6J and **(I)** RAB/6N mice at 2,4, and 7 dpi and in non-infected (mock) mice. **(J)** Semi-quantitative ordinal scoring of lung histopathology in K18-hACE2/6J and RAB/6N mice at 2,4, and 7 dpi, or in non- infected (mock, M) mice. **(K)** Quantification of the percent of CD3+ cells in lungs of K18-hACE2 (black) and RAB/6N (red) mice at 2, 4, and 7 dpi. Quantification was performed on multiplexed immunofluorescent-stained whole lung sections using Halo Image analysis software. Analysis was performed with a Two-way ANOVA. P-values are indicated on the graph. **(L)** Percent lung consolidation in K18-hACE2/6J (black) and RAB/6N (red) mice at 2, 4, and 7 dpi and in non- infected (mock) mice. Quantification was performed on whole lung images using Halo Image analysis software. Analysis was performed with a Two-way ANOVA. P-values are indicated on the graph.

Viral antigen (SARS-CoV-2 N) was observed in both mouse models in alveolar type I and II cells (**Fig. 4D-G**). Antigen staining reflected the increased levels of lung infection in K18- hACE2/6J mice over RAB6/N mice at 2 and 4dpi, and a more gradual antigen dissemination in the latter model prior infection resolution (**Fig. 4D-G**). Histopathological analysis by H&E showed significant lung pathology at 4 and 7 dpi in both models (**Fig. 4H, I, J)**, which was characterized by regional interstitial pneumonia, moderate to severe lymphohistiocytic (albeit with lesser neutrophils in RAB/6N mice), mild to moderate edema, AT2 hyperplasia, and mild single cell necrosis/apoptosis. However, RAB/6N mice exhibited significantly higher levels of T-cell (CD3+) infiltrates at 4 and 7 dpi as well as increased lung consolidation at 7 dpi compared to K18- hACE2/6J mice (**Fig. 4K, L and S8**). Collectively, RAB/6N lung infection is marked by delayed control of viral replication, enhanced T-cell lung infiltration and maladaptive tissue repair.

### Lung infection resolution in RAB6/N mice is associated with increased T-cell extravasation and type-2-based immunity

To define host responses between the two mouse models, we performed flow cytometry analysis on lung extravascular (EV) and intravascular (IV) compartments by intravenously injecting an anti-CD45.2 antibody prior to euthanasia to label circulating immune cells. Analysis was done at 4 dpi (**Fig. S9A-D**), when both models start to exhibit significant lung leukocyte infiltration (**Fig.4H,I,K**).

Consistent with a host response to infection, we observed a significant increase in total CD45+ numbers and EV CD45+ in the lung of both K18-hACE2/6J and RAB/6N upon infection (**Fig. 5A; S10A**). This was further illustrated by significant increases in several activated leukocyte populations in the lungs of both K18-hACE2/6J and RAB/6N compared to mock-infected mice (**Fig. 5B-D; S10A**), including IV CD69+ monocytes, CD11c+ Natural killer (NK) cells and distinct B cell subpopulations, as well as EV CD4+, CD8+ and CD4- CD8- (double negative; dn) T cells, EV CD11c+ dendritic cells (DC), EV NK cells and EV macrophages. Several lineages also elicited a decrease in numbers upon infection, irrespective of the mouse model, such as IV eosinophils and CD11c+ monocytes, and EV conventional DC (cDC), dnT cells/NKT cells, CD11b+CD11c+ DC, and Ly6C- neutrophils (**Fig. S10B,C**). However, despite these similarities, lung IV and EV CD4+ T effector memory (TEM), and naïve EV CD4+ T cells, only increased in RAB/6N mice upon infection, consistent with our histological findings (**Fig. 4K and 5B, C**). EV CD69lo CD8+ T cells cell numbers were also higher in infected RAB/6N compared to K18-hACE2 mice (**Fig. 5C**). Conversely, K18-hACE2/6J mice showed significantly higher numbers of several innate immune cell populations compared to RAB/6N mice at 4 dpi, including EV natural killer (NK) cells and Ly6c+ CD11clo/- CD69+ macrophages (Macrophage – I) (**Fig. 5D**). We also noted a significant increase in EV Ly6CloCD11cintCD69lo macrophages (Macrophage-II) in K18-hACE2/6J mice but not in RAB/6N mice upon infection (**Fig. 5D**). Collectively, our data suggest that RAB/6N and K18-hACE2/6J lung response to SARS-CoV-2 infection are differentially skewed toward T-cell and myeloid responses, respectively.

**Figure 5.**
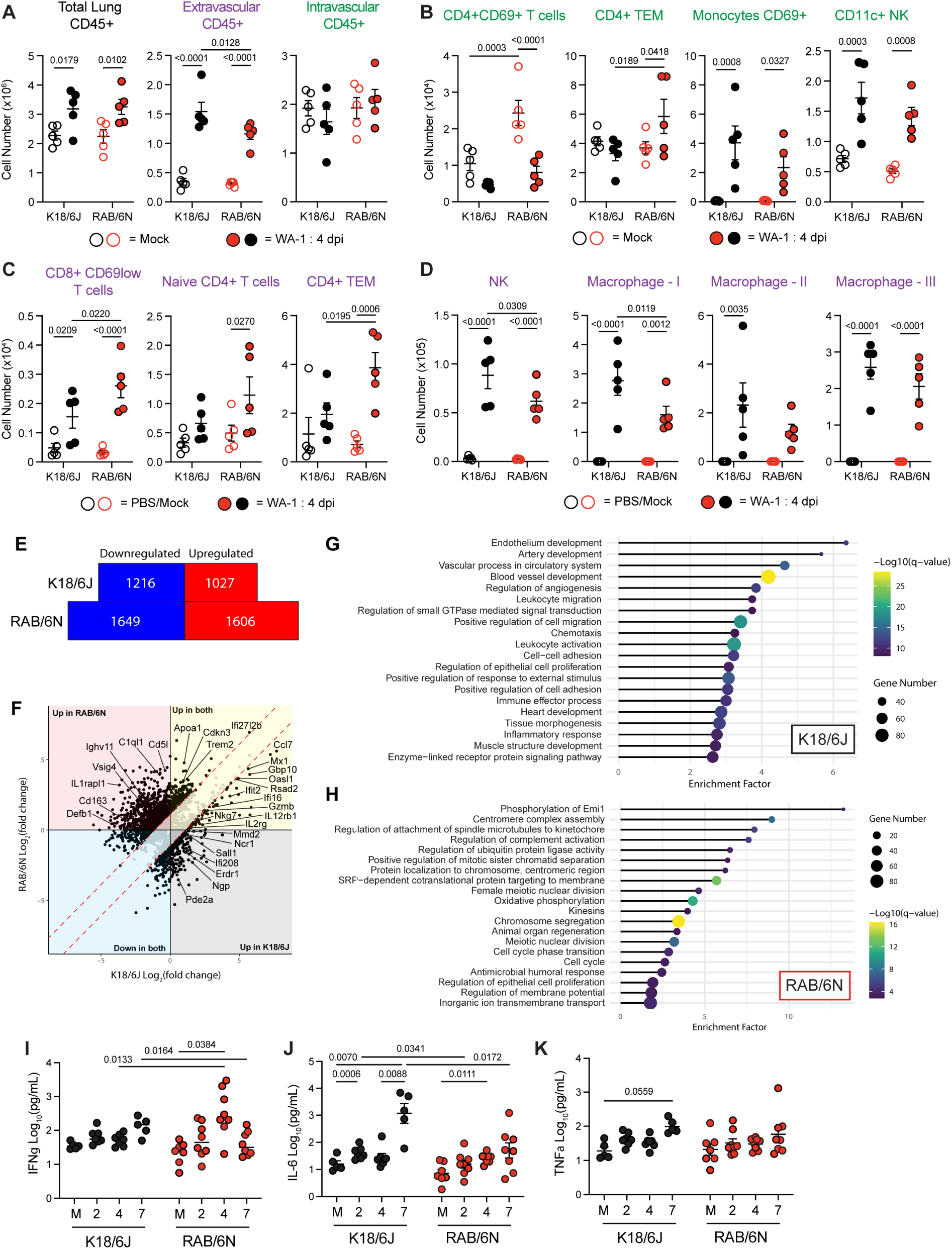
SARS-CoV-2 infection drives differential systemic and lung immune responses during acute infection. (A-D) Flow cytometric analysis of hematopoietic populations in the lungs of K18-hACE2/6J and RAB/6N mice at 4 dpi and in non-infected (mock) mice. Error bars represent mean +/- S.E.M. Analysis was performed using a Two-way ANOVA with Mixed Model effects. P- values are indicated on graphs. **(A)** Total CD45+ cell numbers in the lung (left; black), within the extravascular lung compartment (middle; purple), and within the intravascular lung compartment (right; green). **(B)** Total cell numbers of select immune populations within the intravascular lung compartment. **(C-D)** Total cell number of select immune populations within the extravascular lung compartment. **(E-H)** Transcriptomic analysis (Bulk RNA-sequencing) was performed on lung tissue of K18-ACE2/6J and RAB/6N mice at 4 dpi and in non-infected (mock) mice. **(E)** Total number of downregulated (blue) and upregulated (red) genes in K18-hACE2/6J and RAB/6N mice at 4 dpi and in non-infected (mock) mice. **(F)** Cross plot of differentially expressed genes. Genes up in K18-hACE2/6J and down in RAB/6N are in the black, genes up in both are in yellow, genes down in K18-hACE2/6J and up in RAB/6N are in red, and genes down in both are in blue. **(G, H)** GO-term analysis for genes upregulated in the lungs of K18-hACE2/6J (G) and RAB/6N (H) at 4 dpi. **(I-K)** Serum concentrations of IFNγ **(I)**, IL-6 **(J)**, and TNFα **(K)** in K18-hACE2/6J (black) and RAB/6N (red) mice at 2, 4, and 7 dpi and in non-infected (mock, M) mice. Error bars represent mean +/- S.E.M. Analysis was performed using a Two-way ANOVA with Mixed Model effects. P- values are indicated on graphs.

Bulk RNA-sequencing on whole lung homogenates at 4 dpi revealed 3255 differentially regulated genes in RAB/6N mice and 2243 in K18-hACE2/6J, compared to naïve mice **(Fig. 5E**). While both models elicited a type I IFN-associated response, it was more extensive in K18- hACE2/6J (**Fig.5F, Fig.S11 A, B**). Lungs of K18-hACE2/6J mice were also enriched in pathways related to leukocyte activation, chemotaxis and extravasation while RAB/6N mice were not (**Fig. 5G**). We also observed canonical upregulation of genes associated with macrophage differentiation and activation (*Mmd2*, *Pde2a*, *Ifi208;* along with negative feedback loop genes *Sall1 and Erdr1*) and neutrophil functions (*Ngp*, *Ncr1*) in the lung of this model (**Fig. 5F**). In contrast, transcriptomic signatures in RAB/6N mice suggested enrichment of pathways involved in tissue repair mechanisms (**Fig. 5H**) and the presence of a pro-resolving, M2-like macrophage response (*Cd5l*, *CD163, IL1rapl1, Vsig4)* (**Fig.5F**). Albeit upregulated in both models, several anti-inflammatory and exhaustion markers (*Apoa1*, *Cdkn3*, *Trem2)* were biased toward RABN/6 mice; while pro-inflammatory and type I IFN-associated genes were more biased in K18-hACE2/6J mice (*Ccl7*, *Ifit2*, *Gbp10*, *Mx1* and more; **Fig. 5F**). Notably, RAB/6N mice also simultaneously elicited specific stress and immune signatures associated with complement activation (*C1ql1*) and humoral responses (*Ighv*-associated genes). Together, these findings suggest that while K18- hACE2/6J mice mount canonical immune and inflammatory responses driving lung infection resolution at the peak of viral infection, RAB/6N mice exhibit a type-2, tolerance-biased response trading tissue protection for slower viral clearance.

### RAB6/N and K18-hACE2/6J mount a distinctive systemic inflammatory signature upon acute infection

We then asked whether the differential lung immunological trajectories upon SARS-CoV- 2 infection in the two mouse models correlate with distinctive signatures of systemic inflammatory responses. RAB/6N mice elicited a significant increase in IFNγ in circulation between 0 and 4 dpi, a marker of protective immune responses to SARS-CoV-2 infection(*54–56*). However, no IFNγ response was detected in the blood of K18-hACE2/6J mice (**Fig. 5I**). Rather, IL-6 levels were significantly higher in the serum of K18-hACE2/6J, but not RAB/6N mice, at 7 dpi as compared to naïve mice (**Fig. 5J**). A similar pattern was also observed for peripheral TNFα in K18-hACE2/6J mice, though not significant (p=0.0559) (**Fig. 5K**). These findings underscore that while RAB6/N systemic inflammatory signatures are temporally aligned with lung responses, K18-hACE2/6J systemic signatures are instead temporally synchronized with widespread brain infection.

Collectively, our data suggest that RAB/6N mice could serve as a valuable platform for modeling the systemic impact of SARS-CoV-2 lung infection and inflammation, without the confounding effect of brain infection on systemic inflammation.

### Convalescent RAB/6N mice exhibit systemic inflammation and humoral defects

T-cell-biased host responses, maladaptive lung tissue remodeling, and systemic IFNγ- skewed antiviral responses, a marker of PASC in humans(*38, 57*) suggest a potential for RAB6/N to model PASC. As severe COVID-19 disease and high viral load have been associated with a greater chance of persistence of viral RNA and higher incidence of PASC (*58–60*), we infected RAB/6N mice with 105 PFU of Delta virus, and various tissues were collected from convalescent animals at 30 and 65 dpi (**Fig. S12A**). In total, twenty-five RAB/6N mice (mean survival = 21.2%) across several independently inoculated cohorts were analyzed (**Fig. S12B**). Notably, Delta infection resulted in an increase in peripheral IFNγ levels upon acute infection (**Fig. 6A**), as we reported during WA-1 infection (**Fig. 5I**). Our cohorts of RAB/6N survivors were also slightly biased toward females (27.4% females vs. 14.3% males survived and recovered from Delta infection), consistent with reports that males are more susceptible to fatal COVID-19(*47–49*).

**Figure 6.**
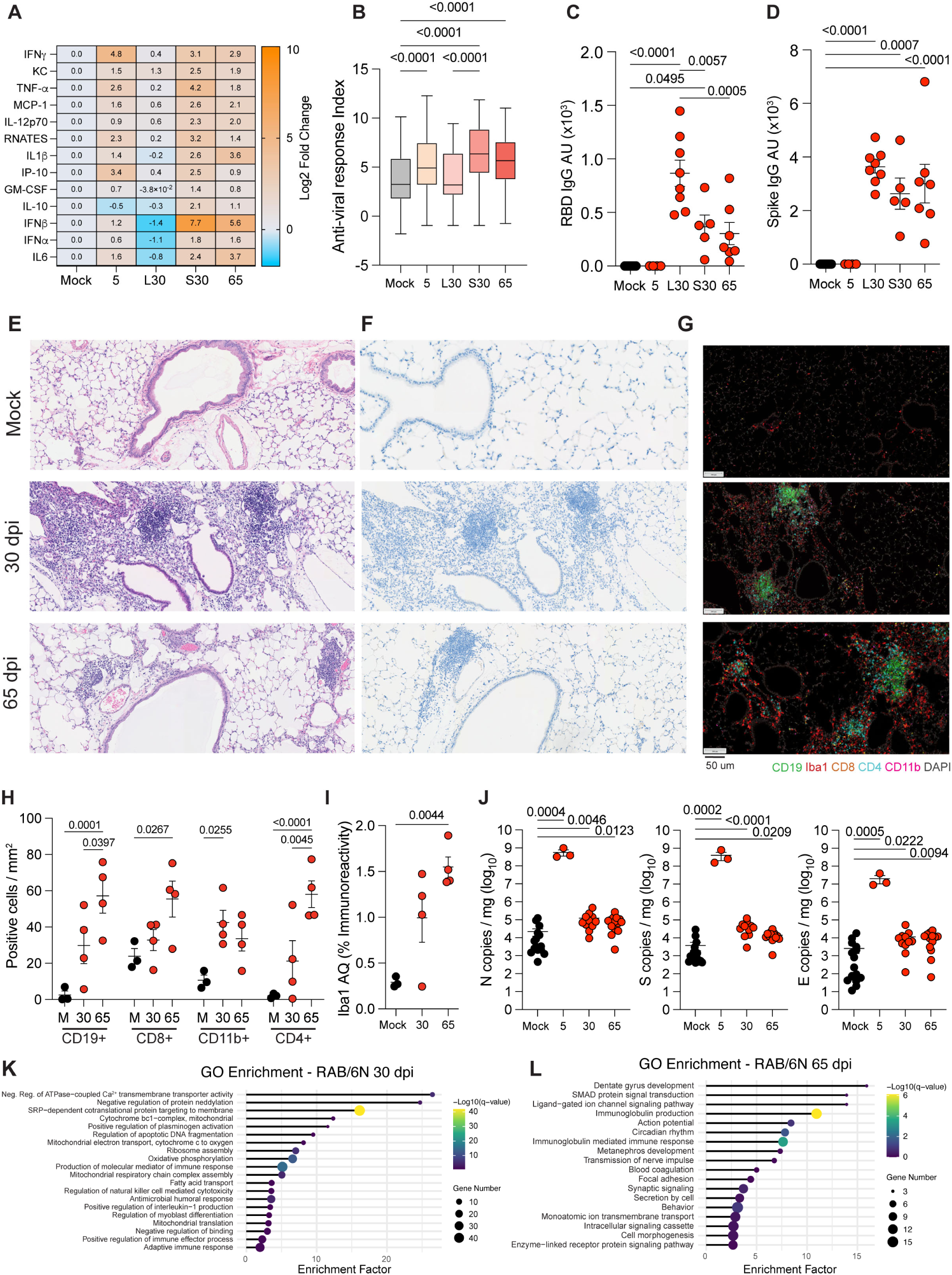
RAB/6N mice display evidence of systemic and pulmonary post-acute sequelae. RAB/6N mice were infected with 1×105 pfu of SARS-CoV-2 Delta variant. Mice surviving infection (∼20%) were characterized at 30 and 65 dpi. **(A)** Mean log2 fold-change of thirteen serum cytokine levels in surviving RAB/6N mice at 30 and 65 dpi compared to non-infected (mock) mice. **(B)** Mean concentration (pg/mL) of thirteen cytokines combined (referred to as anti-viral response index) in surviving RAB/6N mice at 30 and 65 dpi compared to non-infected (mock) mice. For A and B, L = low inflammation and S = severe inflammation at 30 dpi as supported statistically by the anti-viral cytokine index. **(C-D)** SARS-CoV-2 RBD IgG and Spike IgG specific antibody levels in serum of RAB/6N mice at 5 dpi, 30 dpi, and 65 dpi, and in non-infected (mock) mice. L = low inflammation and S = severe inflammation at 30 dpi as determined by cytokine analysis. **(E, F)** H&E **(E)** and SARS-CoV-2 Spike IHC **(F)** on lung tissue sections at 30 and 65 dpi and in non- infected (mock) mice. **(G)** Multiplex IHC (mIHC) on lung tissue sections at 30 and 65 dpi and in non-infected (mock) mice. green = CD19, red = Iba1, orange = CD8, blue = CD4, pink = CD11, and grey = dapi. **(H, I)** Quantification of CD11b+, CD19+, CD4+ and CD8+ cell numbers per mm2 of lung tissue **(H)** and percent of Iba+ cells in analyzed lung tissue section (AQ, area quantification) after mIHC at 30 and 65 dpi and in non-infected (mock, M) mice. **(J)** RT-qPCR analysis of SARS-CoV-2 Envelope (E), Spike (S), and Nucleocapsid (N) genes in lungs of RAB/6N mice at 5, 30, and 65 dpi and in non-infected (mock) mice. **(K, L)** Bulk RNA-sequencing performed on lung tissues of RAB/6N mice at 30 and 65 dpi, and of non-infected (mock) mice. GO-term analysis of significantly upregulated genes at 30 (K) and 65 dpi (L) compared to non- infected (mock) mice. Error bars represent mean +/- S.E.M. Analysis was performed using a Two-way ANOVA with Mixed Model effects. P-values are indicated on graphs.

A defining feature of PASC in patients is long-lasting peripheral inflammation (*61–65*). Thereby, we measured the log2 fold-change of thirteen pro-inflammatory cytokines in the serum of RAB/6N mice at 30 and 65 dpi and compared to naïve RAB/6N mice (**Fig.6A and S13A,B**). At 30 dpi, we noted variable levels of peripheral inflammation with two distinct trajectories: low (L) and severe (S) inflammation (**Fig.6A; S13A,B)**, as defined by the mean concentration (pg/ml) of our thirteen antiviral cytokines combined (referred to as antiviral response index) (**Fig.6B**). However, at 65 dpi, we observed a single inflammatory trajectory marked by aberrantly high levels of multiple inflammatory cytokines, including IFNγ, IL6, IFNβ and IL1β, all of which were reported in excessive amounts in the blood of PASC patients(*57, 65*)(**Fig.6A,B; S13A,B**).

Patients with PASC have also been shown to display dysregulated humoral responses against SARS-CoV-2(*66*). Consistently, we observed a positive correlation between levels of systemic inflammation (mild vs. severe) and serum concentration of anti-SARS-CoV-2 Receptor Binding Domain (RBD) IgG antibodies (**Fig.6C-D**). Of note, concentrations of anti-SARS-CoV-2 spike antibodies were mildly reduced, but not significantly (**Fig.6D**), reflecting a targeted impact of systemic inflammation on RBD-specific antibodies, which typically are more neutralizing, in line with findings from a recent human study(*66*). We refer to RAB/6N mice displaying systemic inflammation as RABPASC mice.

### Lung inflammation and dysfunction signatures in RAB^PASC^ mice

Lungs of RAB6/NPASC mice exhibited the presence of atypical leukocyte aggregates at both 30 and 65 dpi despite the absence of viral antigen (**Fig. 6E-F**). Aggregates were dominantly composed of CD19+ and CD4+ cells, suggesting the presence of cellular structures reminiscent of inducible bronchus-associated lymphoid tissue (iBALT; **Fig. 6G**); whose persistence has been associated with chronic lung inflammation(*67*). Notably, lung tissues experienced a continuous increase in lymphoid aggregates over time (**Fig. 6H**) despite the absence of active viral infection. Further supporting these observations, we observed a significant enrichment of Iba+ cells, a macrophage marker, at 65 dpi but not at 30 dpi (**Fig.6I**). Despite the lack of viral antigen in lung tissues, viral RNA, particularly those coding for S and N proteins, persisted in lung tissues at both 30 and 65 dpi **(Fig. 6J**). These findings are consistent with previous studies identifying persisting viral RNA in PASC patients(*68*).

Bulk RNA sequencing of lung tissues from RAB/6NPASC mice suggested evidence lung inflammation at 30 dpi. This was illustrated by the significant upregulation of genes associated with B cell activation and antibody production (Immunoglobulin light chain: *Igkv12-41, Ighv8-12, Igkv1-88, Igkv4-73*(*69–72*) *and Jchain*(*73*)), stromal and dendritic cell cytokine support (*Flt3L*(*74*)), T-cell proliferation (*G02s*(*75*)) and upregulation of senescence marker (*Gng11*(*76*)) (**Fig. 6K and S14A; Tables 1 and S1**). Pathway enrichment analysis confirmed the presence of a hyperimmune activation state notably defined by enhanced humoral, natural killer cell and IL1β responses, as well as by increased mitochondrial function and metabolism. Reversely, downregulated gene signatures illustrated impaired neutrophil activation and migration (*CD177*(*77*)), dysregulation of epithelial barrier homeostasis (*DMBT1*(*78*)*and Clca3b*(*79*)), defective cilia motility and mucociliary clearance (*Hydin*(*80*)) and decreased goblet cell development and mucus barrier assembly (*Ern2*(*81, 82*)), suggestive of specific lung functional defects that could predispose to multiple lung diseases including allergic airway diseases, interstitial lung diseases, Chronic Obstructive Pulmonary Disease (COPD) and bronchiectasis.

At 65 dpi, lung transcriptomic signatures were enriched for Immunoglobulin(Ig)-coding transcripts **(Fig. 6L and S14B; Tables 1 and S1)**. Of particular interest was *Ighe*, coding for the constant region of IgE, which is a hallmark of type 2 immunity and lung autoinflammation(*83–87*). Upregulation of antimicrobial responses, negative regulators of the mTORC pathway, cell adhesion mechanisms, hypertension and fibrotic mediators, and cancer markers also implicated a wide panel of autoinflammatory disorders and malignancies potentially linked to PASC (**Tables 1 and S1**). Notably, several transcripts also suggested increased crosstalk between the lung and central nervous system (CNS), stressing a functional link between lung inflammation and CNS disorders. We found significant downregulation of several regulators of inflammation, such as the IL-17 pathway, the C1q and TNF-related 4 protein (and phospholipase A2, which have been linked to increased susceptibility to microbial infection(*88–90*) or the development of systemic lupus erythematosus (SLE(*91*)) – an autoinflammatory disorder linked to autoantibody production, such as IgE(*83*) (**Tables 1 and S1**). Loss of *Wnt3* transcripts, and decrease in α-actin-coding, and myosin and myosin-associated genes suggested defects in tissue repair and maintenance mechanisms, with connections to cancers(*92–94*).

### Post-acute infiltration of hematopoietic lineages in RAB^PASC^ mouse tissue is independent of the presence of viral components and associated with neurological disorder signatures

We then aimed to examine the post-acute host responses of tissues not typically recognized as major sites of SARS-CoV-2 replication in humans and that have been associated with PASC manifestations in patients. We first focused on the brain. No statistically significant persistence of viral antigen or RNA was observed in the brain at any time point tested (**Fig. 7A, B**). However, at 30 dpi, the brain exhibited a significant increase in CD3+, CD8+, and GFAP+ cells at 30 dpi compared to naïve animals (**Fig. 7C-E**) despite the absence of viral RNA and viral antigen **(Fig. 7A, B)**. Immune infiltrates returned to naïve levels by 65 dpi (**Fig. 6H,I**). We then leveraged transcriptomic approaches to characterize the tissue-wide impact of this transient hematopoietic infiltration in an unbiased fashion. Bulk RNA-seq on brain tissues at 30 dpi showed a strong signature associated with Ig production, coupled with the activation of potential protective mechanisms against cellular stress and inflammation (**Fig. 7F, G, Tables 1 and S1**). Downregulated gene signatures implied neuroendocrine, brain development, vascular permeability, eosinophil and neuronal transport defects, with implications for neuroinflammatory and neurodegenerative disorders, cancer, cognitive and mood disorders, and increased susceptibility to infection (**Fig. 7F, H, Tables 1 and S1**).

**Figure 7.**
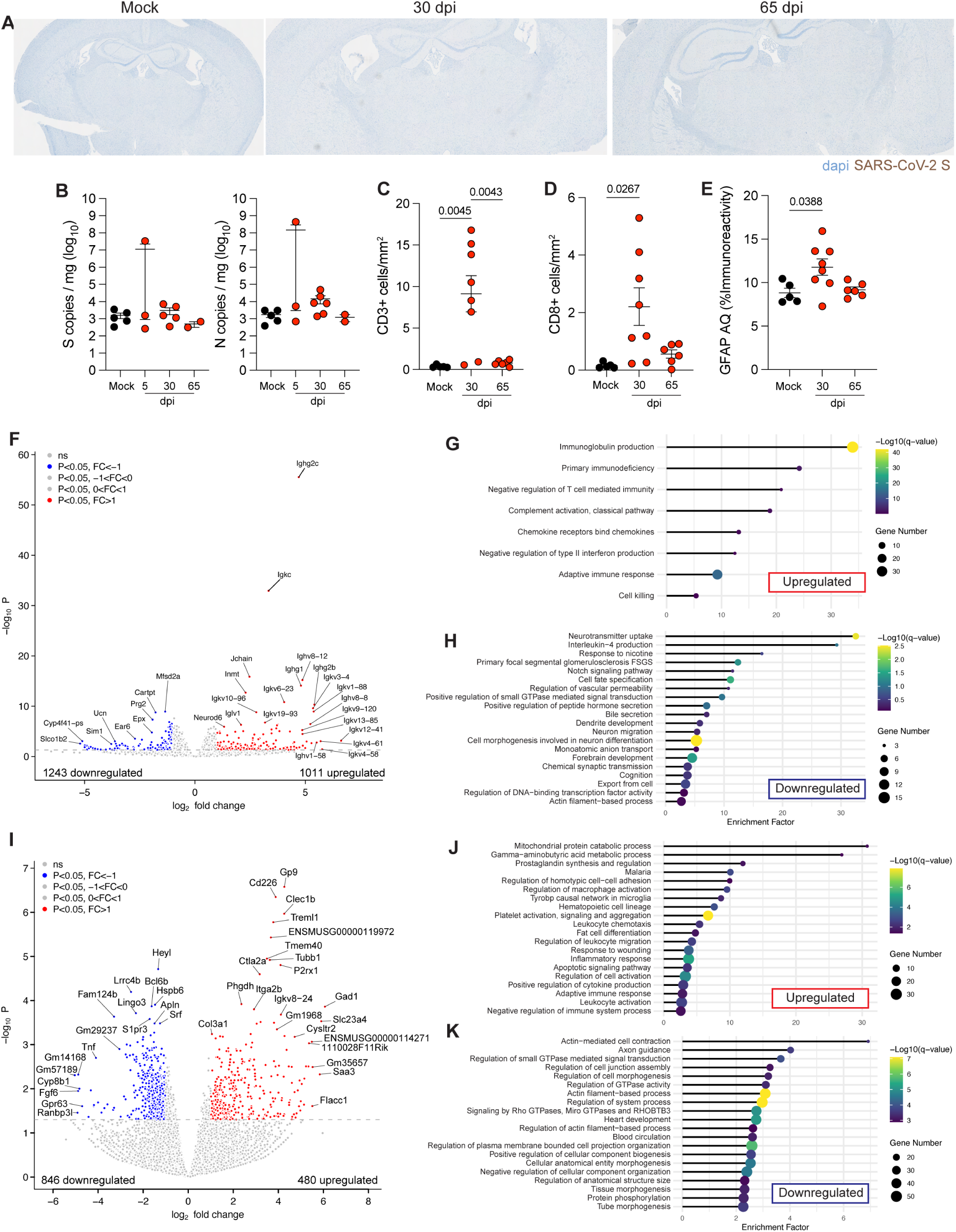
RAB/6N^PASC^ mice display signs of transcriptomic dysregulation in the brain and heart. **(A)** SARS-CoV-2 S IHC was performed on whole brain sections from RAB6NPASC mice at 30 and 65 dpi (1×105 pfu, SARS-CoV-2 Delta virus), as well as for non-infected (mock) mice. Representative images are shown. **(B)** RT-qPCR analysis of Spike (S) and Nucleocapsid (N) genes in the brain of RAB/6N mice at 5, 30, and 65 dpi and in non-infected (mock) mice. **(C-E)** Quantification of CD3+ (C) and CD8+ (D) cell numbers per mm2 of tissue after mIHC, and percent of Iba+ cells (AQ, area quantification) in analyzed tissue sections (E). Two-way ANOVA with Mixed Model effects. P-values are indicated on graphs. **(F-H)** Bulk RNA-sequencing performed on the brain of RAB/6N mice at 30 dpi and of non-infected (mock) mice. **(F)** Volcano plot displaying upregulated (red) and downregulated (blue) genes compared to non-infected (mock) mice. **(G,H)** GO-term analysis of significantly upregulated genes (G) and downregulated genes (H) as compared to non-infected (mock) mice. **(I-K)** Bulk RNA-sequencing performed on the heart of RAB/6N mice at 65 dpi and of non-infected (mock) mice. **(I)** Volcano plot displaying the upregulated (red) and downregulated (blue) genes compared to non-infected (mock) mice. **(J-K)** GO-term analysis of significantly upregulated genes (J) and downregulated genes (K) as compared to non-infected (mock) mice.

Collectively, our findings suggest that post-acute, transient hematopoietic infiltration in the brain can occur weeks following SARS-CoV-2 resolution and in the absence of significant viral RNA or antigen. Such infiltration is associated with a transcriptomic remodeling of the brain with significant implications for the development of chronic neurological disorders.

### Chronic, viral RNA-independent, cardiac inflammation in RAB^PASC^ mice

PASC has been linked to cardiac inflammation and cardiovascular dysfunctions(*64, 95*). While no significant detection of viral RNA was found in the heart at acute (5 dpi) or post-acute (30 and 65 dpi) time points (**Fig. S15**), we found maladaptive transcriptomic signatures at 65 dpi (**Tables 1 and S1**). Upregulated gene signatures suggested evidence of cardiac inflammation in the absence of active viral infection and viral materials (**Fig. 7 I, J**), which was defined by several genes associated with lymphocyte and myeloid activation (**Tables 1 and S1**). We also noted upregulation of genes and pathways related to mitochondrial dysfunction and potential fibrotic remodeling. In parallel, downregulated genes were associated with impaired cardiomyocyte differentiation and integrity, cardiac contractility, vascular repair, cell survival, and protection against oxidative stress, collectively suggesting a dysfunctional or decompensated heart (**Fig. 7I, K, Tables 1 and S1**).

Together, our findings suggest that the heart can exhibit chronic inflammation despite the absence of acute infection and presence of viral RNA in RAB/6N mice. This inflammation is defined by signatures of cardiac inflammation and dysfunction, which have been associated with an increased risk of cardiac failure and cardiometabolic diseases.

## DISCUSSION

Our study reports RAB/6N mice as a novel hACE2 transgenic mouse model susceptible to SARS-CoV-2 clinical isolates and recapitulating several of the systemic and multifaceted features observed in human PASC cases. RAB/6N mice elicit severe and potentially fatal disease upon SARS-CoV-2 infection, albeit they display a lesser susceptibility to fatal outcomes compared to K18-hACE2/6J mice in a variant and viral dose-specific manner. Lethal outcomes can manifest without any evidence of brain infection, and when observed, neurodissemination remains minimal and regionalized. The most notable differences between the RAB/6N and K18-hACE2/6J mice are found within their lung and systemic antiviral responses. RAB/6N mice exhibit delayed initiation of viral clearance, which is associated with enhanced T-cell extravasation in the lung, type-2 tolerance-biased immune and maladaptive lung consolidation, as compared to a canonical type-1 and myeloid-biased immune response in K18-hACE2/6J mice. Systemically, RAB/6N mice elicit an increase in IFNγ levels that temporally aligns with pulmonary infection, whereas K18- hACE2/6J mice exhibit an increase in IL-6 and TFNα, which is concomitant with widespread brain infection. Adult convalescent RAB/6N mice elicit systemic inflammatory signatures in the peripheral blood up to two months post-infection and persistence of viral RNA in lung tissues, which is consistent with findings from PASC patients(*61–65, 68*). Notably, tissue inflammation was independent of the presence of persisting viral RNA, as it was observed both in the lungs as well as in other peripheral tissues free of viral RNA. Leveraging RAB/6NPASC mice, we also uncovered tissue-specific, molecular correlates of various disorders and pathologies linked, or merely implied, to PASC in humans, ranging from autoimmune diseases and cardiovascular issues to cancers (**Fig. 8**; **Table 1 and S1**). Together, our work illuminates this model as a promising platform to dissect the mechanisms underpinning SARS-CoV-2 acute and post-acute pathogenesis, including how acute maladaptive antiviral responses may seed post-acute sequelae, as well as to evaluate PASC therapies.

**Figure 8.**
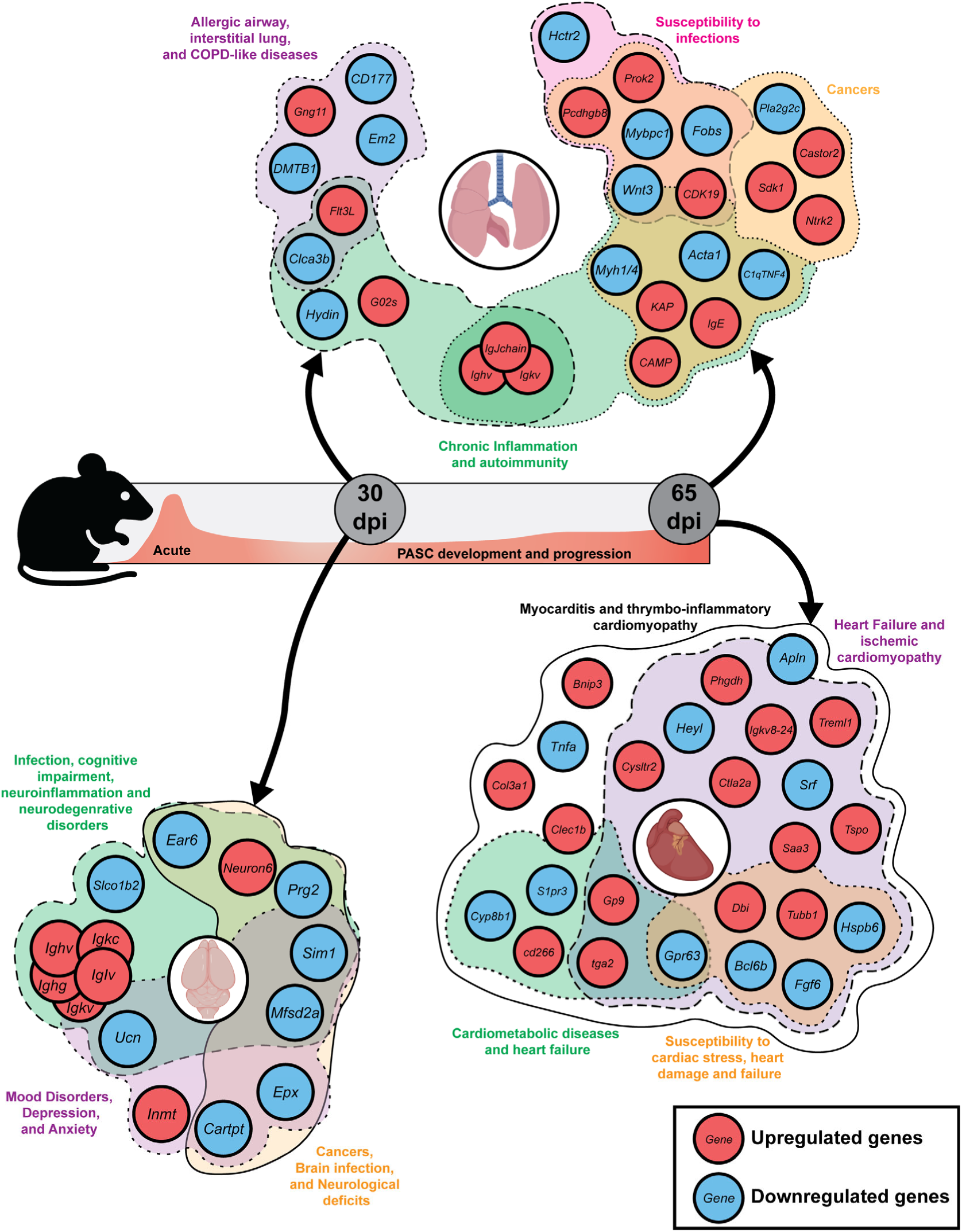
Gene network map of dysregulated disease pathways in RAB/6N^PASC^ tissues. Schematic showing select significantly upregulated (red) and downregulated (blue) genes in the lungs and brains of RAB/6N mice at 30 dpi and in the lungs and hearts at 65 dpi, which are linked to various disease pathways. Created with BioRender.com.

**Table 1.**
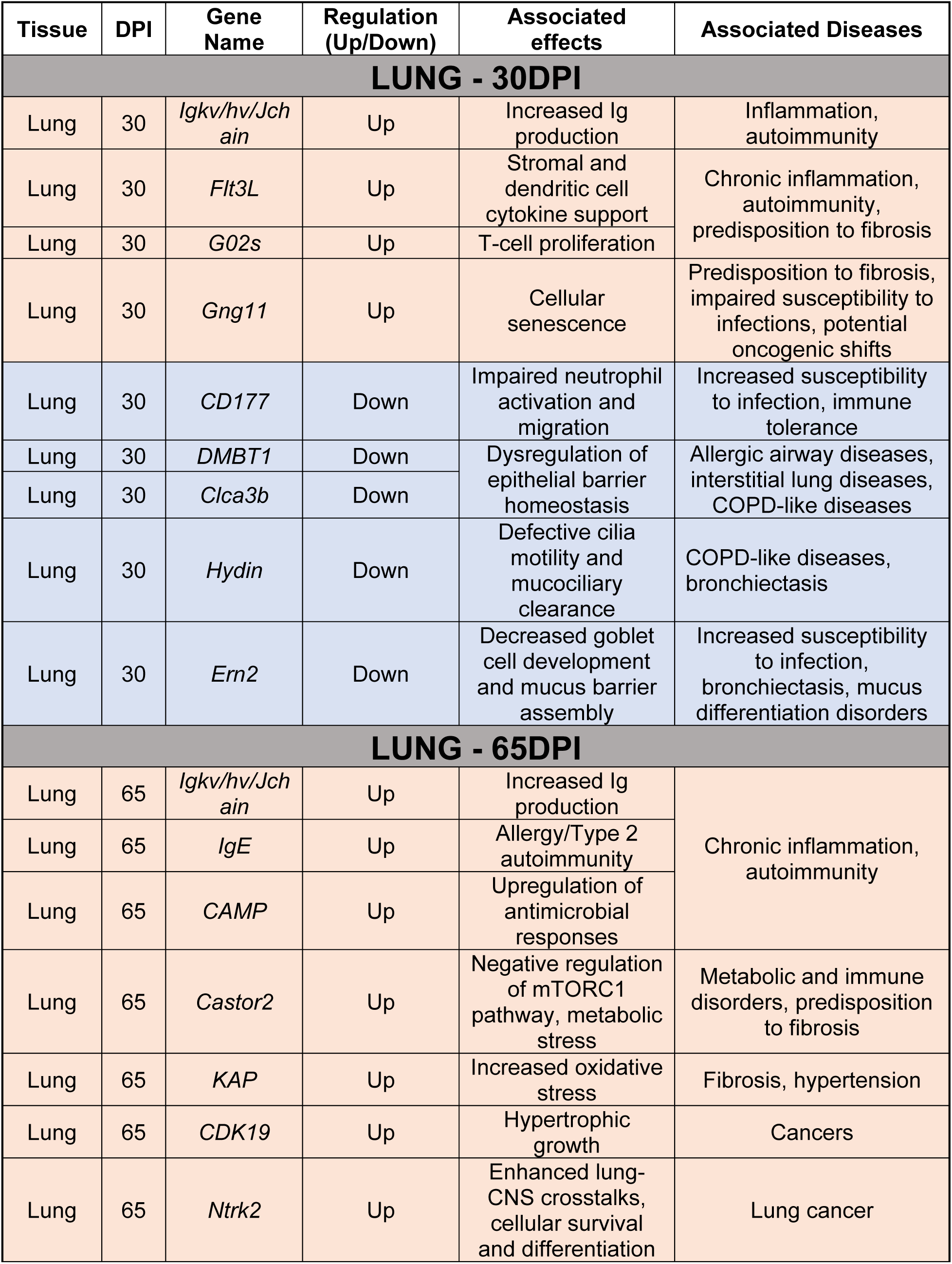
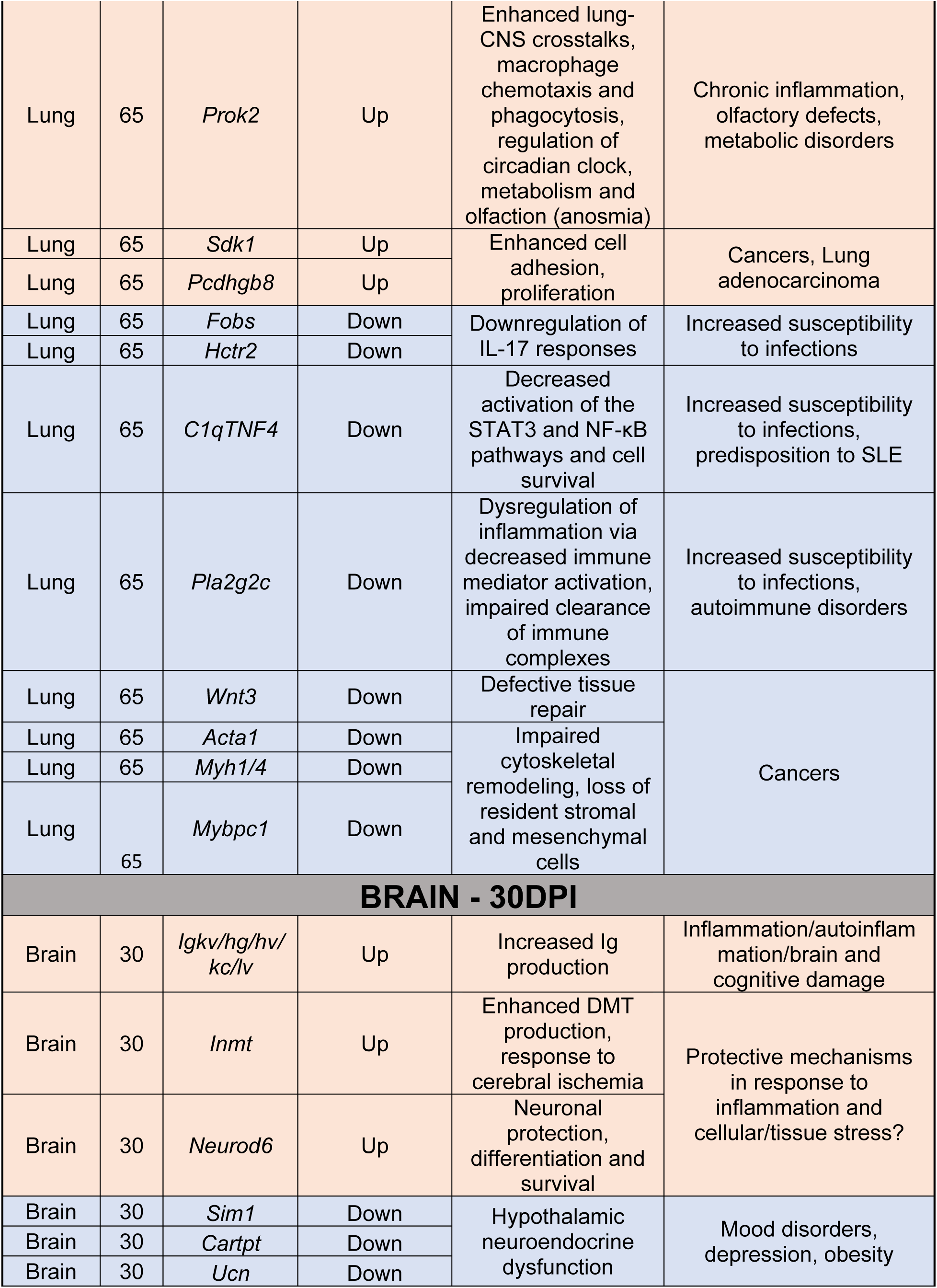
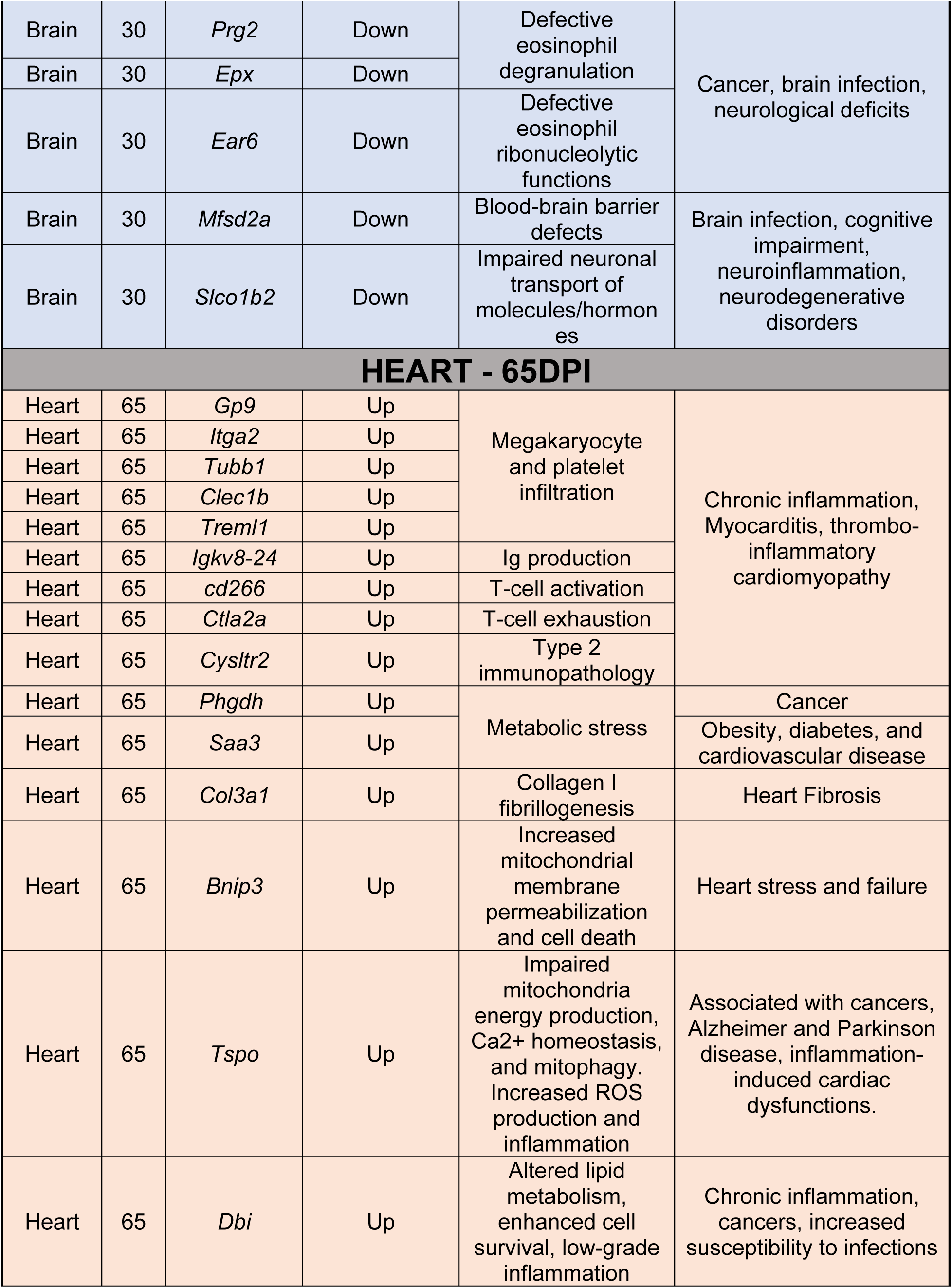
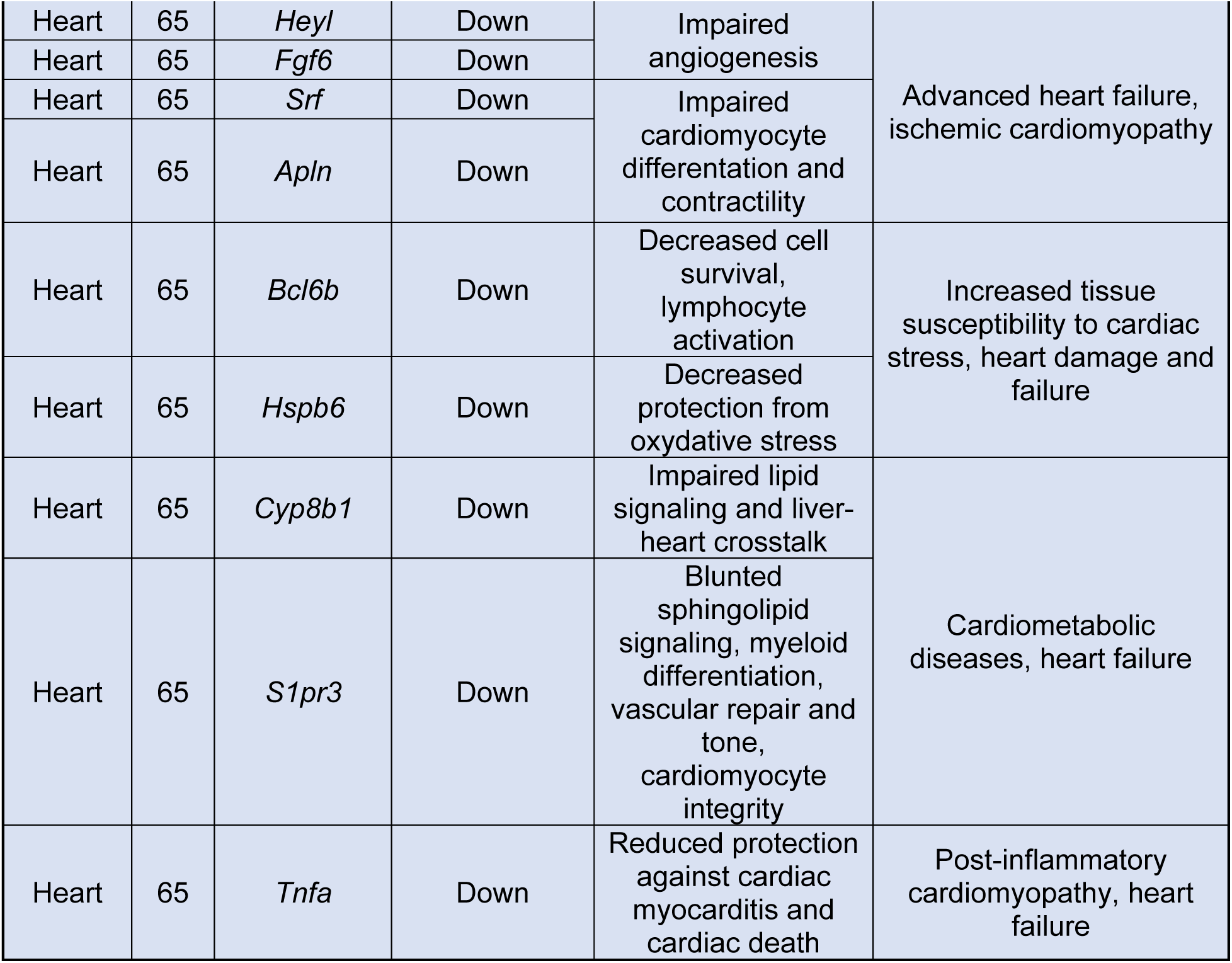
List of select genes significantly up- (light red) or downregulated (light blue) in the lung, brain and heart of RAB/6N mice at 30 and/or 65 dpi. Associated effects and disease phenotypes are listed for each gene showing statistically significant differential regulation. Literature references for each gene and associated effect/diseases are available in Supplemental Table 1.

Despite RAB/6N mice having a single copy of the K18-hACE2 cassette per cell, lung *hACE2* expression was not significantly different compared to K18-hACE2/6J mice. This observation suggests the potential importance of the C57BL/6N sub-strain background in driving the phenotype of RAB/6N mice compared to the K18-hACE2/6J mouse model. C57BL/6J mice carry a deletion in the *Nnt* gene (*42–44*), as well as a missense mutation in the *Nlrp12* gene associated with a neutrophil recruitment defect(*96*). Conversely, C57BL/6N have a deletion and point mutation in the *Crb1* and *Cyfip2* genes, respectively(*97, 98*). Defective expression of *Nnt* and *Nlrp12* in C57BL/6J mice suggests that inflammatory and stress responses may be differentially regulated in this sub-background compared to C57BL/6N mice, consistent with literature reports(*96, 99, 100*). An interesting focus of future investigations will be to determine whether the genetic features of the 6N background directly drive the systemic acute and post-acute immune responses and inflammation following SARS-CoV-2 exposure, or whether these responses are mostly the result of a 6N-dependent differential control of early viral replication in the upper and/or lower respiratory tract.

Another feature of the RAB/6N mouse model is its partial susceptibility to brain infection, along with limited neurodissemination when brain infection occurs. Notably, these features align more closely with human and other animal model studies compared to the abnormal widespread brain infection observed in K18-hACE2/6J mice. Further research will be required to establish whether these phenotypes are caused by specific inflammatory responses in the olfactory bulb and/or the brain of RAB/6N mice. In any case, our findings highlight RAB/6N mice as a platform to capture mechanisms governing susceptibility to SARS-CoV-2 neuroinvasion (leveraging the model’s partial susceptibility to this event) as well as the mechanisms that may limit neurodissemination (through comparison with the K18-hACE2/6J model). By extension, this also calls for further investigation into delineating the causes of lethality in RAB/6N mice. While our data support evidence for enhanced maladaptive lung responses to infection in RAB/6N mice, whether these pulmonary features are the major driver of death will have to be determined.

Clinical manifestations of PASC vary greatly, ranging from excessive fatigue and brain fog to heart conditions and pulmonary dysfunctions(*34, 35, 101*). PASC has been associated with systemic and persistent inflammation(*61-65*), dysregulated innate, cellular and humoral responses(*61, 63, 66, 102*), iron dysregulation(*95*), and coagulation defects(*95*), highlighting the multifaceted nature of this/these disease(s). While patient studies suggest that persistent immune activation(*61-65*), residual viral materials(*68, 103, 104*), or dysregulation of major endocrine pathways(*105, 106*) may drive specific features of PASC, these studies remain descriptive and are not amenable to dissecting the underlying mechanisms driving PASC or its pathophysiological consequences. For these reasons, several animal models of PASC have been developed over the past few years, from non-human primates to specific mouse models(*36*). One animal model that has been suggested to be useful for our understanding of PASC relies on using MA strains of SARS-CoV-2 (MA-PASC model). This model can recapitulate persistent lung sequelae consistent with pulmonary PASC, including lung inflammation and fibrosis in aged C57BL/6J or BALB/c mice(*20, 37–39, 107*). It has enabled the identification of multiple molecular and cellular mediators of pulmonary PASC, including an IFNγ-associated pro-fibrotic monocyte-derived macrophage response(*38*) and persisting CD8+-mediated stimulation of lung macrophages, which chronically release IL1β in a peroxisome-dependent manner(*37, 107*). Though the MA- PASC model is a robust model of pulmonary sequelae, its overreliance on aged animals, the lack of systemic disease and inflammation, and the absence of persistence of viral materials have limited its applications. Our findings demonstrate that RAB/6N mice can model systemic PASC in adult mice and the multifaceted nature of this disease using clinical isolates of SARS-CoV-2. The ability to trigger PASC in young to middle-aged animals also makes this model valuable for studying age-related effects of PASC and more cost-effective by removing the need to age animals for PASC studies.

Our findings from this study suggest that pulmonary type-2-biased and systemic IFNγ responses may be important contributors to RAB/6N’s ability to recapitulate systemic inflammatory syndromes and disease profiles consistent with PASC patients. Future investigations will be valuable in investigating how the characterized acute antiviral responses may be variant-specific and how they can be translated into the development of distinct PASC profiles. This line of thought also directly implies that the superior ability of RAB/6N mice to recover from severe disease, as well as their reduced susceptibility to brain infection, may not be critical determinants for modeling PASC, but rather valuable additions providing logistical and bias-mitigating benefits to the model. Consistently, recent studies have reported the inability of convalescent K18-hACE2/6J mice to model robust and systemic PASC phenotypes(*40*).

Notably, RAB/6N mice recapitulate many systemic PASC features reported in human patients that have not been reported in previous animal models. This includes the persistence of systemic pro-inflammatory cytokines in the peripheral blood(*62, 64, 65*), reduced antibody titers(*66*), persistence of viral RNA in active replication sites(*68, 104*) as well as signatures of cognitive impairment(*108*), blood-brain barrier defect(*62*), cardiac inflammation and dysfunction(*64*), and signatures of autoimmunity(*109, 110*) (**Fig. 8**, **Table 1**). Our model also recapitulates evidence of lung inflammation, increased predisposition to fibrosis and lung epithelial dysfunction signatures, as with the MA-PASC model(*37–39, 107*). The absence of significant lung fibrosis in RABPASC mice could suggest that this is an age-dependent phenotype in mice, consistent with findings from the MA-PASC model(*20, 38, 39*). Our PASC findings in RABPASC mice go beyond patient and mouse studies by unraveling the impact of systemic PASC on the molecular signatures of many tissues, from the lung to the heart. We expose transcriptomic regulations and networks of genes defining known PASC-associated disease phenotypes, such as lung dysfunctions, myocarditis and cognitive impairments and many other diseases postulated to be linked to PASC, such as cancers, cardiometabolic diseases, neurodegenerative diseases and specific autoimmune disorders (e.g., SLE). A notable feature of RABPASC mice is their robust Ig upregulation signature in the lung and brain. This signature is in potential alignment with reports proposing a causative link between autoantibodies and PASC symptoms(*111, 112*), and could suggest the potential RABPASC mice to replicate autoantibody-induced PASC disorders; a hypothesis that will require further investigation. Another recent study unraveled that acute SARS-CoV-2 infection can increase cancer-associated deaths in mouse models and individuals predisposed to cancers(*113*). Whether PASC can predispose to cancer is a distinct question that would have to be addressed. While the transcriptomic heart signatures of patients previously infected with SARS-CoV-2 and affected with myocarditis have been characterized(*114*), the PASC diagnosis of these patients and a causal association between their previous infection exposure and myocarditis were not established. Our work provides evidence of a link between PASC and cardiac inflammation, allowing for future studies into the mechanisms driving cardiac disease. Together, our findings suggest that PASC could seed or predispose to a wide range of chronic and debilitating human diseases, warranting future experiments to functionally evaluate the incidence of these diseases at the cellular, tissue and organism levels in RAB/6N mice. These lines of investigation also open opportunities to assess the impact of extrinsic and intrinsic factors, including comorbidities, diet, and age, in driving the incidence and/or severity of these PASC-associated diseases.

As the drivers of PASC-associated systemic inflammation remain unknown, whether persisting viral RNA (or other residual viral components) contributes to this process has been a matter of debate. Our findings suggest that there is no positive association between tissue inflammation and the presence of viral RNA. However, local persistence of viral RNA, such as in the lungs, could sustain proximal inflammatory responses that diffuse systemically in peripheral tissues free of viral components. RAB/6N mice offer a unique opportunity to assess how persistent viral RNA contributes to PASC-associated systemic inflammation, to mechanistically dissect how it may do so, and to assess the benefits of PASC therapeutic strategies targeting residual viral materials. The transcriptomic resources in this study also represent an initial blueprint of the different disease ramifications of PASC (**Fig. 8**; **Table 1 and S1**), which could be expanded and validated in future studies to anticipate the long-term clinical management of PASC. More generally, this model also represents an *in vivo* platform to dissect how maladaptive acute immune responses and infection-induced chronic inflammation can drive the emergence of various disease processes, from malignancies to hematopoietic dysfunctions. Beyond SARS-CoV-2 and PASC, our findings also suggest that RAB/6N mice may stand as a valuable platform to model maladaptive acute immune responses and post-acute diseases in the context of many other hACE2-dependent βCoV, including specific bat βCoV with pandemic potential.

### Limitations of the study

Our study has several limitations. First, as the K18 promoter drives hACE2 expression, the expression remains contained in the epithelial compartment and not in other specific immune cells, such as macrophages. Notably, this suggests that abortive macrophage infection is not a critical driver of systemic PASC, and that improving RAB/6N mice to recapitulate macrophage abortive infection could further refine our ability to model PASC in this mouse model. An additional limitation routinely associated with mouse work is the immunologically naïve nature of our mouse model, precluding our ability to assess how previous infection and inflammatory stimuli, notably via trained immunity, may regulate the development of physiologically relevant PASC phenotype – a line of investigation of significant interest for future experiments. While the limited survival of RAB/6N mice upon severe SARS-CoV-2 disease requires large cohort sizes to achieve sufficient statistical power for PASC studies, the unique characteristics of the RAB/6N mice over existing models likely outweigh this limitation. However, a comprehensive characterization of the drivers of death in these animals upon SARS-CoV-2 infection and the use of PASC-causing variants associated with less lethal outcomes (i.e., high-dose Omicron variant infection), will provide avenues to further enhance the power of RAB/6N mice for PASC studies.

## ACKNOWLEDGMENTS

This work was supported in part by a start-up fund and Peter Paul Career Development Professorship from Boston University (to F.D.), Clinical and Translational Science Awards (grant UL1 TR001430) from the National Center for Advancing Translational Sciences of the National Institutes of Health (to F.D. and N.A.C.) and a Massachusetts Consortium on Pathogenesis Readiness (MassCPR) grant (to R.B.C.). D.K. received support from an NIH T32 Immunology Training Program (T32AI007309) and Hartwell Fellowship awarded by The Hartwell Foundation (55212221). M.C. received support from an Institutional Development Award (IDeA) from the National Institute of General Medical Sciences of the National Institutes of Health (grant P20GM130555-5011), U.S. Department of Agriculture’s (USDA) National Institute of Food and Agriculture (NIFA) Agriculture and Food Research Initiative and American Rescue Plan Act through USDA Animal and Plant Health Inspection Service (APHIS) competitive grant number 2023-70432-39465 and the School of Veterinary Medicine, Louisiana State University (PG009641). The content is solely the responsibility of the authors and does not necessarily represent the official views of the National Institutes of Health. We thank the Evans Center for Interdisciplinary Biomedical Research at Boston University Chobanian & Avedisian School of Medicine for their support of the Affinity Research Collaborative on ‘Respiratory Viruses: A Focus on COVID-19’. This work utilized a Ventana Discovery Ultra and Vectra Polaris that were purchased with funding from National Institutes of Health SIG grants (S10 OD026983 & S10OD030269). This work was also supported by the Boston University Flow Cytometry Core Facility. We thank the Boston University Animal Science Center, and the NEIDL animal core staff for their outstanding support. We also thank all the Douam, Crossland, Connor, Taconic Team, NEIDL members, and members of the department of virology, immunology and microbiology and pathology and laboratory medicine biochemistry at Boston University for their constant support and advice. Some figures were created using Biorender.com. We used Grammarly (Grammarly Inc.) to assist with grammar correction and sentence structure optimization during manuscript preparation.

## AUTHOR CONTRIBUTIONS

D.K. and F.D. designed the overall study framework. D.K., G.U., N.A.C. and F.D. conceptualized the study. D.K., G.U., A.E.T., N.A.C. and F.D. designed experiments and key assays. D.K., A.E.T., G.U., M.M., A.K.O., C.MC., S.G., F.F-S, and F.D. performed experiments. M.M. and J.N. maintained the mouse colonies and generated animals for experiments. A.E.T., A.O.C., M.C., M.L., H.P.G. and N.A.C. performed histological and immunohistochemistry/hybridization assays and supported data analysis. J.L. and C.H. carried out computational analysis. A.K., N.P, and K.P.F. supported bioluminescence imaging analysis. J.W. and K.A. generated the RAB/6N mice. J.E.S-C, A.B. and R.B.C. provided access to key resources. A.K., N.P, K.P.F., A.B., J.W., K.A., R.B.C. and C.H. provided conceptual and technical inputs and/or helped with data interpretation. D.K. and F.D. wrote the manuscript with contributions from all authors.

## DECLARATION OF INTERESTS

D.K., G.U., A.E.T., J.W., K.A., N.A.C. and F.D. are co-inventors on a U.S. provisional patent application (Serial No. 63/832,794) related to the application of RAB/6N mice to investigate acute and post-acute SARS-CoV-2 pathogenesis. K.P.F. reports that he was, at the time of this study, an employee of PerkinElmer, Inc., a manufacturer of diagnostic and analytical equipment. N.P. and A.K. declare the following competing interests as shareholders of InVivo Analytics with issued patents. The rest of the authors declare no conflict of interest.

## MATERIALS AND METHODS

Detailed descriptions of the materials and methods are provided in the Supplemental Information.

### Study design

The goal of this study was to develop and characterize a novel hACE2 transgenic mouse model (RAB/6N) to better model acute and post-acute COVID-19 disease. To generate this model, a cDNA cassette harboring the hACE2 gene under the control of hK18 promoter was integrated through a targeted insertion into the ROSA26 locus of C56BL/6NTac mice. 12-30 weeks old male and female hemizygous RAB/6N mice, along with the widely used K18-hACE2 mice (which incorporates a similar hACE2 expression cassette), were then infected with various SARS-CoV-2 variants at different doses through the intranasal route or with 1xPBS (mock), and longitudinal disease progression and survival were determined. For comprehensive acute infection studies, we compared the dynamics of clinical disease, viral replication in the lung and brain, and antiviral responses between hemizygous RAB/6N mice and K18-hACE2 mice following WA-1 inoculation (105 PFU). To do so, we leveraged bulk-RNA sequencing, multiplex flow cytometry, histopathological analysis, multiplex cytokine analysis, *in vivo* imaging technologies and virological assays. For post-acute infection studies, hemizygous RAB/6N mice were infected with Delta (105 PFU) and tissues from convalescent animals were collected at 30 and 65 dpi. Bulk-RNA sequencing, multiplex IHC and cytokine analysis, ELISA assays, and RT-qPCR were used to characterize evidence of post-acute inflammatory syndromes, tissue dysfunctions, hematopoietic defects, and viral persistence. For all experiments except RNA sequencing and large-panel flow cytometry analysis, two to three independent experiments were performed with mice ranging from 12-30 weeks.

### Institutional approvals

All animal experiments described in this study were performed in accordance with protocols that were reviewed and approved by the Institutional Animal Care and Use and Committee of Boston University (PROTO202000020; Approval date: 06/15/2020). All mice were maintained in facilities accredited by the Association for the Assessment and Accreditation of Laboratory Animal Care (AAALAC). All replication-competent SARS-CoV-2 experiments were performed in a biosafety level 3 laboratory (BSL-3) at the Boston University National Emerging Infectious Diseases Laboratories (NEIDL).

### Data availability

Raw data from our Bulk-seq analyses will be made available upon publication of the manuscript.

### Statistical Analysis

For all survival curve comparisons, a Log Rank (Mantel-Cox) test was performed with p-values of <0.05 being statically significant. For viral quantification (pfu), histology quantification, flow cytometry, cytokine, IgG quantification, and RT-qPCR analysis was performed using an One-way or Two-way analysis of variance (ANOVA) test as described. All error bars are indicated as mean ± standard error of the mean (SEM). All statistical tests and graphical depictions of results were performed using GraphPad Prism version 10.1.1 software (GraphPad Software, La Jolla, CA). All p-values are indicated on graphs.

