## Supplemental Information for "A Mouse Model of SARS-CoV-2-Driven Acute Maladaptive Responses and Chronic Systemic Diseases"

### FOR

**CONTENT:**

**1. Supplemental Text**

**Supplemental Text 1: Mouse models of SARS-CoV-2 infection**

**Supplemental Text 2: The importance of mouse genetic background**

**2. Supplemental Materials and Methods**

**3. Supplemental Figures and Legends (8 Supplemental Figures)**

**Figure S1. Schematic showing the strategy used to generate the targeted transgenic ACE2 knock-in allele in RAB/6N mice.**

**Figure S2. K18-hACE2/6J and RAB/6N show distinct lung transcriptomes at baseline.**

**Figure S3. Male and female weight loss dynamics after SARS-CoV-2 challenge.**

**Figure S4. Correlation plot of dose and percent lethality in RAB/6N and K18-hACE2/6J mice.**

**Figure S5. Omicron BA.1 infection causes sex-biased disease.**

**Figure S6. Quantification of neuroinvasion in RAB/6N mice.**

**Figure S7. *In vivo* imaging of RAB/6N mice reveals sustained infection.**

**Figure S8. Halo quantification of immune populations from multiplex immunohistochemistry.**

**Figure S9. Unbiased immune profiling of the lung immune environment by flow cytometry.**

**Figure S10. Immune populations in the lung that trend similarly in K18-hAEC2/6J and RAB/6N mice.**

**Figure S11. K18-hACE2/6J and RAB/6N mice show robust transcriptional responses in the lung at 4 days post-infection.**

**Figure S12. Development of PASC model from mice surviving infection with Delta.**

**Figure S13. RAB/6N$^{PASC}$ mice show persistent systemic inflammation.**

**Figure S14. RAB/6N PASC mice show signs of multi-tissue gene dysregulation.**

**Figure S15. RAB/6N$^{PASC}$ mice are free of viral RNA in heart tissues**

**4. Supplemental Table**

**Supplemental Table 1. List of select genes differentially regulated in RAB/6N$^{PASC}$ mice with associated literature references.**

**5. Supplemental References**

#### SUPPLEMENTAL TEXT

##### Supplemental Text 1: Mouse models of SARS-CoV-2 infection

Genetic humanization of the mouse ACE2 locus does not render mice susceptible to clinical disease upon SARS-CoV-2 infection(*1*). However, random insertion of one or several copies of the hACE2 cDNA into the mouse genome, either under the control of the K18 or CMV enhancer/chicken beta-actin (CAG) promoter, confers hypersusceptibility to pulmonary SARS-CoV-2 infection and fatal brain infection(*2-7*). In contrast, insertion of a single copy of the K18-hACE2 cassette into the mouse collagen COL1A1 locus results in clinical disease but no brain infection(*8*). However, the lack of significant lung inflammatory responses in this model during acute infection underscores the need for further investigation to establish its suitability for studying COVID-19 immunopathology, as well as PASC.

##### Supplemental Text 2: The importance of mouse genetic background

Beyond the impact of transgene integration strategy, mouse genetic background can also regulate susceptibility to SARS-CoV-2 infection(*9-12*). Notably, sub-background can also matter as C57BL/6J but not C57BL/6N mice lack the mitochondrial antioxidant enzyme, Nicotinamide nucleotide transhydrogenase (Nnt)(*13-15*), which affects mitochondrial function and glucose metabolism(*16*) – two processes that closely regulate both immune responses(*17*) and SARS-CoV-2 infection(*18*). The functional impact of such differences has been illustrated by the higher resistance of C57BL/6JN mice to inflammation-associated disease upon influenza A virus infection(*19*), or the lesser lung inflammatory response to dsRNA of C57BL/6N mice(*20*), as compared to C57BL/6J mice.  As hACE2-transgenic mouse models built on the C57BL/6N background and expressing hACE2 under a ubiquitous promoter (i.e., CAG) have not shown any benefits over K18-hACE2-6/J mice (built on the C57BL/6J background), including a lesser susceptibility to viral neuroinvasion(*6, 7*), we hypothesized in our study that targeted insertion into a safe harbor locus of a C57BL6/N strain (C57BL6/NTac) of the hACE2 cDNA under the control of the epithelial promoter could model distinctive infection and pathogenesis course as compared to other hACE2-transgenic mouse models upon SARS-CoV-2 infection.

#### METHODS

##### Mouse Strains

RAB/6N mice were developed in collaboration with Taconic Biosciences (see "Generation of RAB/6N mice" below) for this study. These animals are commercially available as ROSA26-K18-hACE2 mice (Item 18675) from Taconic Biosciences. Hemizygous B6.Cg-Tg(K18-ACE2)2Prlmn/J (K18-hACE2/6J) mice were purchased from Jackson Laboratories (Bar Harbor, ME, USA) (strain # 034860). Animals were group-housed by sex in Tecniplast green line individually ventilated cages (Tecniplast, Buguggiate, Italy). Mice were maintained on a 12:12 light cycle at 30–70% humidity and provided ad-libitum water and standard chow diets (LabDiet, St. Louis, MO, USA).

##### Cell lines

VeroE6 cells were grown in Dulbecco's modified Eagle's medium (DMEM) (ThermoScientific, Waltham, MA, USA) supplemented with 5-10% (v/v) heat-inactivated fetal bovine serum (FBS) (Bio-Techne, R&D systems, Minneapolis, MN, USA) and 1% (v/v) penicillin streptomycin (P/S) (ThermoScientific, Waltham, MA, USA). Caco2 cells expressing human ACE2 and human TMPRSS2 were kindly provided by the laboratory of Dr. Mohsan Saeed at Boston University. Caco-2 hACE2/hTMPRSS2 cells were cultured in DMEM with 5-10% FBS, 1% P/S, 2.5 ug/mL of puromycin and 2.5 ug/mL of Blasticidine. All cell lines were maintained in a cell incubator at 37°C with 5% $CO_2$.

##### SARS-CoV-2 Virus Stocks

All replication-competent SARS-CoV-2 experiments were performed in a BSL-3 facility at the Boston University NEIDL. The clinical isolate named 2019-nCoV/USA-WA1/2020 strain (NCBI accession number: MN985325) of SARS-CoV-2 was obtained from BEI Resources (Manassas, VA, USA). The SARS-CoV-2 B.1.617.2 (Delta) and Omicron BA.1 isolate was kindly provided by the laboratory of Dr. John H. Connor at Boston University. Recombinant SARS-CoV-2 virus expressing a NanoLuciferase reporter (rSARS-CoV-2 NL) in place of ORF7b within the WA-1 strain [46] was generously provided by the Laboratory of Dr. Pei-Yong Shi, University of Texas Medical Branch (UTMB). SARS-CoV-2 WA-1, Delta and WA-1/NanoLuc viruses were grown and titrated on VeroE6 cells and Omicron BA.1 was grown and titrated on Caco2 hACE2/hTMPRSS2 cells. For growth, $8x10^6$ cells were plated into a T-175 flask the day prior to infections. The next day cells were infected with a multiplicity of infection (moi) = 0.1 pfu/cell of passage 1 (P1) of WA-1, Delta, WA-1/NanoLuc, or BA.1 in 10 mL of Opti-MEM media + GlutaMax (Gibco®, Carlsbad, CA, USA) and incubated for 1 hr at 37°C to allow for viral adsorption. After 1 hr incubation, 10 mL of DMEM with 4% FBS was added to cells, incubated for 24 hr, and then media was replaced with 25 mL of DMEM with 2% FBS and incubated at 37°C. Cells were assessed for cytopathic effect (CPE) and virus was collected when significant CPE was observed (between 60-96 hr post infection).

Cell supernatant was clarified by centrifugation for 10 min at 350 *x g* at 4°C and filtered through 0.22 uM filter. WA-1, Delta and WA-1/NanoLuc Virus was overlayed over a 20% sucrose cushion (Sigma-Aldrich, St. Louis, MO, USA) and concentrated by ultracentrifugation (Beckman Coulter Optima L-100k; SW32 Ti rotor) at 25,000 RPM for 2 hours at 4°C. Viral pellets were then suspended in 1xPBS titrated by plaque assay on Vero E6 cells (see *Viral quantification by plaque assay*). For Omicron BA.1, virus was mixed at a 1:1 ratio with PEG6000 (Sigma-Aldrich, cat# 1546580) and incubated at 4°C overnight, centrifuged at 12,000 *x g* for 30 min at 4°C, then suspended in 1xPBS.

#### Generation of RAB/6N (ROSA26-K18-hACE2-C57BL/6N) mice

The RAB/6N mouse line was generated by recombinase-mediated cassette exchange (RMCE). Human ACE2 was synthesized according to NCBI transcript NM_021804.3 and cloned downstream of the KRT18 5' region (K18EpilongSEAP) and a translational enhancer (TE)(*21*). The 3' region of KRT18 (K18i6x7pA)(*22*) was inserted downstream of ACE2. This construct was cloned into an RMCE vector carrying F3 and FRT sites for site specific integration and a promoter-less NeoR selection marker. The final vector and a Flp-expression plasmid were co-transfected into a C57BL/6NTac embryonic stem cell (ESC) line containing an F3/FRT-RMCE landing pad in the ROSA26 locus. Recombinant clones resistant to neomycin treatment were picked and expanded for analysis. Extensive molecular validation by Southern Blotting, sequencing and quantitative PCR ensured correct recombination and absence of random integrations or aneuploidy. Validated ESC-clones were injected into BALB/c blastocysts for chimera generation. Chimerism was assessed by coat color. Sperm of several highly chimeric animals was analyzed for relative content of the targeted allele and overall quality after cryopreservation. Selected sperm was then used in *in vitro* fertilization of C57BL/6NTac oocytes to generate hemizygous RAB/6N G1-animals. RAB/6N mice were then maintained by back-cross with C57BL/6NTac wild-type mice.

#### SARS-CoV-2 infections in mice

Male and female 12-30-week-old hemizygous K18-hACE2/6J and RAB/6N mice were intranasally inoculated with SARS-CoV-2 in 50 µL of sterile 1xPBS split between both nostrils. Non-infected mice (mock) were inoculated with 50 uL of sterile 1xPBS. All infections were performed while mice were under anesthesia with 1-3% isoflurane. After inoculations mice were observed until they were awake from anesthesia.

#### Clinical scoring and monitoring for survival

For survival studies, mice were monitored for changes in body weight, altered respiration, general changes in appearance, responsiveness and neurological signs of disease. An IACUC-approved clinical-scoring system was used to monitor disease progression and define humane endpoints. The score of “1” was given for each of the following situations: body weight, 10–29% loss; respiration, rapid and shallow with increased effort; appearance, ruffled fur and/or hunched posture; responsiveness, low to moderate unresponsiveness; and neurological signs, tremors. The sum of these individual scores constituted the final clinical score (0-5). Mice were considered moribund and humanly euthanized in case of weight loss greater than or equal to 30%, or if they received a clinical score of 4 or above for two consecutive days. Body weight and clinical score were recorded once per day for the duration of the study. For the purpose of survival curves, mice euthanized on a given day were counted dead the day after. Mice that were found dead in the cage were counted dead on the same day. For euthanasia, an overdose of ketamine/xylazine was administered followed by a secondary method of euthanasia.

**Tissue collection**

At predetermined endpoints for tissue collection (2, 4, 5, 7, 30 and 65 dpi) or at human euthanasia criteria, mice were euthanized by ketamine/xylazine overdose. Tissues were then collected and placed in either 1) 10% Formalin for at least 72 hrs for viral inactivation prior to histopathological analysis or 2) 600-750 uL of RNALater (MilliporeSigma: #R0901500ML) and stored at -80C for downstream analysis.

***In vivo* 3D-imaging and analysis**

Before imaging, mice were administrated two, 75 µL subcutaneous injections of 1X PBS containing 0.65 uM Fluorofurimazine (FFz) substrate (Promega, Madison, WI, USA) for a total of 1.3 uM FFz per mouse. Mice were then imaged using a 3D-imaging mirror gantry isolation chamber (InVivo Analytics, New York, NY, USA) and an IVIS spectrum imager (PerkinElmer,

Waltham, MA, USA). To perform 3D imaging, mice were anesthetized with 2.5% isoflurane, placed into a body conforming animal mold (BCAM) and InvivoPLOT™ mirror gantry (InVivo Analytics), and then imaged within 5 min of FFz injection. Images were acquired using a sequence imaging as followed; 60 s (s) open filter, 240 s 600 nm, 60 s open, 240 s 620 nm, 60 s open, 240 s 640 nm, 60 s open, 240 s 660 nm, 60 s open, 680 nm, 60 s open. Data analysis was performed using the cloud based *In Vivo* Plot software (In Vivo Analytics).

**RNA extraction from tissue**

For RNA extraction, 20–30 mg of tissue was weighed out, placed into a 2 mL tube with 600 µL of RLT buffer with 1% β–mercaptoethanol and a 5 mm stainless steel bead (Qiagen, Hilden, Germany: #69989), then homogenized using a Qiagen TissueLyser II (Qiagen) with the following cycle: two min dissociation at 1800 oscillations/min, one min rest, two min dissociation at 1800 oscillations/min. Homogenized tissues were centrifuged at 13,000 rpm for 10 min at room temperature and cleared supernatant was transferred to a new 1.5 mL eppendorf tube. RNA extractions were performed using a Qiagen RNeasy Plus Mini Kit (Qiagen: #74134), according to the manufacturer's instructions (Qiagen: #79256). RNA was eluted in 30 µL of RNase/DNase free water and quantified by Nanodrop2000.

**Tissue processing for viral quantification**

30-50 mg of tissue was weighed out and placed into a 2 mL tube with a 5 mm stainless steel bead and 600 uL of OptiMEM. Tissues were homogenized using a Qiagen TissueLyser II (Qiagen) with the following cycle: two min dissociation at 1800 oscillations/min, one min rest, two min dissociation at 1800 oscillations/min. Dissociated tissues were centrifuged at 13,000 rpm for 10 min at room temperature and cleared supernatant was transferred to a new 1.5 mL eppendorf tube for downstream analysis.

**Viral Quantification by plaque assay**

WA-1, Delta, and WA-1/NanoLuc were all quantified by plaque assay on VeroE6 cells. Omciron BA.1 was quantified by plaque assay on Caco2 hACE2/hTMPRSS2 cells. For plaque assay, serial dilutions of concentrated and non-concentrated stocks were generated and then 300 uL of each dilution was plated onto 12-well plates and incubated at 37°C with 5% $CO_2$ for 1 hour, followed by the addition of 1 mL of 1.2% Avicel in DMEM containing 2% FBS and 1% Penn/Strep. Cells were incubated for 3-days before the overlay was removed; cells were then fixed with 10% neutral buffered formalin and stained with crystal violet (0.1% crystal violet in 10% ethanol/water).

**SARS-CoV-2 E-specific reverse transcription quantitative polymerase chain reaction (RT-qPCR)**

Viral RNA was quantitated using single-step RT-quantitative real-time PCR (Quanta qScript One-Step RT-qPCR Kit, QuantaBio, Beverly, MA, USA; VWR; #76047-082) with primers and TaqMan® probes targeting the SARS-CoV-2 E gene as previously described(*23*). The 20 µL reaction mixture contained 10 µL of Quanta qScript™ XLT One-Step RT-qPCR ToughMix, 0.5 µM of each primer E_Sarbeco_F1 (5'-ACAGGTACGTTAATAGTTAATAGCGT-3') and E_Sarbeco_R2 (5'-ATATTGCAGCAGTACGCACACA-3'), 0.25 µM of FAM-BHQ1 probe E_Sarbeco_P1 (FAM-ACACTAGCCATCCTTACTGCGCTTCG-BHQ 1), and 2 µL of template RNA. RT-qPCR was performed using an Applied Biosystems QuantStudio 3 (ThermoFisher Scientific) and the following cycling conditions: reverse transcription for 10 min at 55 °C, an activation step at 94 °C for 3 min followed by 45 cycles of denaturation at 94 °C for 15 s and combined annealing/extension at 58 °C for 30 s. For absolute quantitation of viral RNA, a 389 bp fragment from the SARS-CoV-2 E gene was cloned onto pIDTBlue plasmid under an SP6 promoter using NEB PCR cloning kit (New England Biosciences, Ipswich, MA, USA). The cloned fragment was then in vitro transcribed (mMessage mMachine SP6 transcription kit; ThermoFisher) to generate an RT-qPCR standard.

#### SARS-CoV-2 N and S reverse transcription quantitative polymerase chain reaction (RT-qPCR)

Viral RNA was quantitated using single-step RT-quantitative real-time PCR according to manufacturer's instructions (Luna® Universal One-Step RT-qPCR Kit, kit #E3005L, New England Biolabs) with primers targeting the SARS-CoV-2 N and S genes; S_FW (5'-AATTTAGTGCGTGATCTCCC -3') and S_RV (5'- AACTTCTATGTAAAGCAAGTAAAG-3'), N_FW (5'- CCAGAATGGAGAACGCAGT-3') and N_RV (5'-TGAGAGCGGTGAACCAAGA-3'). RT-qPCR was performed using Applied Biosystems QuantStudio 3 (ThermoFisher Scientific) following Luna® Universal One-Step RT-qPCR Kit manufacturer's instructions. For absolute quantitation of viral RNA, SARS-CoV2 Delta RNA was extracted with *Quick*-RNA Viral Kit (Zymo Research, kit #R1035) from 1ml of viral stock to generate an RT-qPCR standard. Standard viral RNA copies were calculated according to the REF protocol following RNA quantification with NanoDrop2000.

#### RT-qPCR analysis of *hACE2*

RT-qPCR was performed using the Luna universal one step RT-qpCR kit #E3005L, New England Biolabs). The manufacturer's protocol was followed with minor adjustments for a final volume of 12 µL consisting of 2 µL of RNA, 2 µL of forward primer, 2 µL of reverse primer, 0.6 µL of 20X Luna WarmStart RT enzyme mix and 6 µL of 2X Luna universal one-step reaction mix. Using the Applied Biosystems QuantStudio 3 (ThermoFisher Scientific), the 12 µL reaction mixture used the following cycling conditions: 50ºC for 10 minutes (reverse transcription), 95ºC for 2 minutes (initial denaturation) followed by 40 additional cycles of 95ºC for 15 seconds (denaturation) and 60ºC for 1 minute (annealing/extension) and finally a melt curve analysis. The melt curve analysis began with a 30 second waiting period at 65ºC with temperature change of +0.5ºC/s and a 5 second pause after every increment during which the machine acquired fluorescence at each new temperature until it reached 95ºC. Threshold cycle values (Ct) also known as quantification cycle (Cq) were determined using QuantStudio Design and Analysis software V1.5.1. Normalization was performed using mouse *Gapdh* as a control. Primers are as followed: h*ACE2* forward primer: 5'-CGAAGCCGAAGACCTGTTCTA-3' and reverse primer: 5- GGGCAAGTGTGGACTGTTCC-3'; m*Gapdh* forward primer 5'-GGTGCTGAGTATGTCGTGGAGTCTA-3' and reverse primer 5'-AAAGTTGTCATGGATGACCTTGG-3'.

**Cytokine analysis using BioLegend LegendPlex**

Cytokine analysis was performed on serum that was inactivated using a 1:1 dilution in 2% Tween-80 prepared in 1x PBS for a final concentration of 1% Tween-80. Serum-Tween-80 mixtures were incubated at 4°C for 2 hours to allow for inactivation and then analyzed using a LEGENDplex™ Mouse Anti-Virus Response Panel (13-plex) (Biolegend, cat # 740622) kit as per the manufactures protocol. This kit test for the following cytokines: IFN-γ, CXCL1 (KC), TNF-α, CCL2 (MCP-1), IL-12p70, CCL5 (RANTES), IL-1β, CXCL10 (IP-10), GM-CSF, IL-10, IFN-β, IFN-α, IL-6. Sample collection was performed using an BD Biosciences LSRII Flow cytometer and data was analyzed using BioLegend Browser based "Data Analysis Software Suite for LegendPLEX" (https://legendplex.qognit.com/user/login?next=home).

**SARS-CoV-2 antiviral antibody measurements**.

Antibodies reactive to SARS-CoV-2 RBD and Spike were assayed from plasma using a previously reported ELISA assay(*24*). Briefly, wells of 96-well plates (Thermo Fisher, cat# 34028) were coated with 50μL/well of 2μg/mL or 1μg/mL SARS-CoV-2 RBD (a gift from the Schmidt lab at the Ragon Institute) or 1μg/mL SARS-CoV-2 Spike (R&D Systems, cat# 10549-cv-100,) in sterile PBS for one hour at room temperature. Following incubation, the coating solution was removed, and the plate was washed three times with 200μL of sterile PBS and then blocked with 200μL of casein blocking solution (Thermo Fisher, cat# 37528) for one hour at room temperature. Following blocking, the plate was washed again as described. Plasma samples, diluted 1:50 or 1:100 in casein, were added to the plate at 50µL/well. Monoclonal SARS-CoV-2 RBD reactive antibodies (IgG, clone 4, Sino Biological, cat#40591-MM41) were also diluted in casein and added to the plate at 50µL/well. Dilution buffer alone was added to blank wells and the plate was incubated for one hour at room temperature. Diluted samples were then removed, and the plate was washed three times with 0.05% PBS-Tween 20 (0.05% PBST). Immediately after, anti-mouse horseradish peroxidase-conjugated secondary antibodies for IgG (Thermo Fisher, cat# 31430) diluted 1:5000 in casein were added to all wells at 50µL/well and incubated for 30 minutes at room temperature. Next, the plates were washed four times with 0.05% PBST, and then 50µL of 3,3',5,5'-Tetramethylbenzidine (TMB)-ELISA substrate solution (Thermo Fisher, cat# 34028) was added to all wells, and incubated in the dark. Incubation occurred until a difference in color between the seventh dilution (~1.37ng/mL) and diluent-only 'zero' well was visualized (8-10 minutes). Once a difference was observed, ELISA stopping solution for TMB was added at 50µL/well and optical density was measured at 45 nm (OD 450 nm) on a SpectraMax190 Microplate Reader.

**Determination of Arbitrary Units**

Data was analyzed using GraphPad Prism 10. Arbitrary Units (AU) on a ng/mL scale were calculated from logarithmic transformation of the optical density (OD) values after subtraction of averaged blank (diluent only) wells. Non-linear regression of the sigmoidal standard curves, generated by known amounts of monoclonal anti-SARS-CoV-2 RBD IgG, was used to extrapolate a "concentration" for each sample, and then inverse log-transformed and multiplied by the dilution factor. The linear portion of the dilution curve for antigen-coated well, adjusted for by subtraction of paired diluent-only values, was used to determine the net AU value per sample.

**Tissue Processing for flow cytometry analysis**

For flow cytometry analysis, 5-8 minutes prior to euthanasia, mice were retro-orbitally injected with 2ug of anti-CD45.2 BUV737 (clone: 104, BD Biosciences, cat # 612778) to allow for antibody labeling of immune cells in circulation. Then, mice were euthanized, and lungs were stored in RPMI with 2% FBS prior to single-cell suspension generation using a "Lung Dissociation Kit, mouse" (Miltenyi Biotec, Auburn, CA, USA; cat # 130-095-927) per the manufacturer's protocol. Briefly, lungs were placed into gentleMACS™ C Tubes (cat # 130-093-237) containing 2.4 mL of 1X Buffer S, 100 uL of Enzyme D and 15 uL of Enzyme D. Tissues were subject to dissociation using a gentleMACS Dissociator with program m_Lung_01, placed at 37°C for 30 minutes with constant rotation/agitation, then dissociated a second time using a gentleMACS Dissociator with program m_Lung_02. After dissociation, cells were filtered through a 70uM filter, centrifuged at ~300 *x g* for 8 minutes at 4°C, supernatant was removed and cells were suspended in 3 mL of ACK lysis buffer (Gibco, cat # A1049201). Cells were then incubated for 5 minutes at room temperature and lysis of red blood cells was quenched with 10 mL of RPMI with 2% FBS. Cells were then counted and used for downstream flow cytometry staining.

**Staining cells for flow cytometry analysis**

After generation of single cell suspension, $10x10^6$ - $20x10^6$ cells were used for flow cytometry staining. Cells were centrifuged at 300 *X g* for 8 min at 4°C. The cell pellet was resuspended in 50 uL Zombie NearIR (Biolgend, cat# 423106, 1:800 dilution) live/dead staining in 1x PBS and incubated for 20 minutes at on ice. Cells were then washed twice with 200 uL of BD stain buffer (BD biosciences, cat# 554656). After washing cells were incubated with 25 uL of BD stain buffer with 1 uL of TruStain FcX (anti-mouse CD16/32 antibody) (Biolegend, cat# 101320) and 2.5 uL True-Monocyte blocking buffer (Biolegend, cat# 426103) and incubated for 10 min at room temperature. After blocking, 25 µL of antibody cocktail containing antibodies and 10uL BrilliantViolet Staining buffer (BD Biosciences, cat# 563974) was added to each sample and incubated in the dark for 30 min at 4°C. After staining, 200 uL of BD stain buffer was added to each sample, samples were centrifuged at 300 *x g* for 8 min at 4°C, washed with 250 uL BD stain buffer, centrifuged at 300 *x g* for 8 min at 4°C, and then fixed in 200 µL 4% PFA for 1 hour. After fixation cells were washed twice with 250 uL PBS, resuspended in PBS, and stored protected from light at 4°C until analysis.

**Flow cytometry Data Acquisition and Analysis**

Flow cytometry analysis was performed using a Cytek Aurora spectral analyzer (Cytek Biosciences). SpectraFlo (Cytek) software was utilized for spectral unmixing of the data using ordinary least square algorithm. Unmixed flow cytometry data were analyzed using OMIQ software from Dotmatics (www.omiq.ai, www.dotmatics.com). Data processing pipeline was assembled within the Omiq.ai cloud computation platform (Dotmatics). Live $CD45^{+}CD45.2^{+}$ (lung intravascular) and live $CD45^{+}$ $CD45.2^{-}$ (lung extravascular) immune cells were concatenated, asinh transformed (cofactor = 6000) and clustered with Phenograph algorithm (k = 30, distance metric = euclidean)(*25*) and projected into opt-SNE(*26*) space (perplexity = 30, theta = 0.5, opt-SNE endpoint = 5000; PCA pre-initialization embedding). Clusters were annotated based on phenotypes, color-coded and overlaid on the opt-SNE projection graphs. Each marker Mean Fluorescence Intensities (MFIs) of clustered datasets were organized into hierarchically clustered heatmaps.

**Antibodies for flow cytometry**

Anti-CD11b BUV395 (clone: M1/70, BD Biosciences, cat #: 563553) 1:200, anti-CD45.2 BUV737 for i.v. (clone: 104, BD Biosciences, cat #: 612778) 2ug/mouse, anti-NK11 BV421 (clone: PK136, Biolegend, cat#: 108731) 1:50, anti-Ly-6C eFluor450 (clone: HK1.4, Invitrogen, cat# 48-5932-82) 1:200, anti-CD45 BV510 (clone: 30-F11, Biolegend, cat#: 103138) 1:200, anti-CD44 BV570 (clone: IM7, Biolegend, cat#: 103037) 1:50, anti-CD19 BV605 (clone: 6D5, Biolegend, cat#: 115539) 1:200, anti-CD62L BV650 (clone: MEL-14., BD Biosciences, cat#: 564108) 1:800, anti-

CD43 BV750 (clone: S7, BD Biosciences, cat# 747277) 1:200, anti-CD11a BV786 (clone: M17, BD Biosciences, cat# 740866)1:200, anti-CD8 AF488 (clone: 53-6.7, Biolegend, cat# 100723) 1:800, anti-CD64 FITC (clone: X54-5/7.1, Biolegend, cat# 139316) 1:200, anti-B220 Spark Blue 550 (clone: RA3-6B2, Biolegend, cat# 103265) 1:50, anti-CD4 PerCP (clone:GK1.5, Biolegend, cat# 100431) 1:50, anti-I-A/I-E (MHC-II) PerCP Cy5.5 (clone: M5/114.15.2, BD Biosciences, cat# 562363) 1:200, anti-CD69 PE (clone: H1.2F3, Biolegend, cat# 104508) 1:100, anti-PD-1 PE Dazzle594 (clone: RMP1-30, Biolegend, cat# 109115) 1:50, anti-F4/80 PE-Fire 640 (clone: QA17A29 , Biolegend, cat#: 157319) 1:50, anti- CD11c PE Cy7 (clone: HL3, BD Biosciences, cat# 558079) 1:100, anti-CD103 APC (clone: 2E7, Biolegend, cat# 121414) 1:50, anti-CD3 Alexa647 (clone: 145-2C11, Biolegend, cat# 100322) 1:400, anti-Ly6G Alexa 700 (clone: 1A8, BD Biosciences, cat# 561236) 1:200, and anti-Siglec-F APC Cy7 (clone: E50-2440, BD Biosciences, cat# 565527) 1:200.

**Bulk RNA-sequencing analysis**

Adapters and low-quality bases were removed from raw reads using Trimmomatic (0.39)(*27*), remaining sequences were mapped to the mouse genome (GRCm39.107) using STAR (2.7.10b)(*28*) and a count table was produced using subread (2.0.1)(*29*). Differentially expressed genes analysis was performed with R (4.1.3)(*30*) using DESeq2 (1.34.0)(*31*). For each comparison, genes with pval $< 0.05$ and absolute value of log2 fold change $> 1$ were considered significant and separated into upregulated and downregulated for over representation analysis using Metascape(*32*).

**Tissue Preparation for Histological Analysis**

Following euthanasia, lungs were inflated with 1% low-melt agarose (Fisher Scientific, Cat# BP165-25) through the trachea to preserve alveolar architecture prior to fixation. All tissues, including lung and brain, were immersed in 10% neutral-buffered formalin (NBF) for a minimum of 72 hours to ensure complete viral inactivation before removal from the BSL-3 facility. Brains were sectioned coronally using a mouse brain matrix (Zivic Instruments). Fixed tissues were processed using a Tissue-Tek VIP-5 vacuum infiltration processor (Sakura Finetek USA, Torrance, CA, USA) and embedded in paraffin with a HistoCore Arcadia embedding station (Leica Biosystems, Wetzlar, Germany). Formalin-fixed, paraffin-embedded (FFPE) blocks were cut into 5 µm sections using an RM2255 rotary microtome (Leica), and sections were mounted onto positively charged glass slides (Fisherbrand Superfrost Plus) for staining.

**Histology and Brightfield Immunohistochemistry**

Hematoxylin and eosin (H&E) staining was performed using a Leica Autostainer XL (Leica Biosystems, Wetzlar, Germany). Brightfield immunohistochemistry (IHC) was performed on a Ventana Discovery Ultra autostainer (Roche, Basel, Switzerland). For detection of viral antigen, slides were incubated with either monoclonal rabbit anti-SARS-CoV-2 spike (S1) antibody (clone E5S3V; Cell Signaling Technology, Cat# 99423; 1:400 dilution) or mouse monoclonal anti-SARS-CoV-1/2 nucleocapsid protein antibody (clone 1C7C7; Cell Signaling Technology, Cat# 68344; 1:1000 dilution). Antigen retrieval for S1 protein was performed using CC1 buffer (Tris-based, Roche Cat# 950-500124) at 95 °C for 1 hour, followed by primary antibody incubation at 37 °C for 32 minutes. For nucleocapsid, CC2 buffer (citrate-based, Roche Cat# 950-223) was used for 32 minutes at 91 °C, followed by incubation at room temperature for 40 minutes. Detection was performed using ChromoMap DAB Detection Kit (Roche, Cat# 760-159), with hematoxylin counterstaining and coverslipping. Pathologic evaluation was performed qualitatively and semi-quantitatively by a board-certified veterinary pathologist (N.A.C.).

**Multiplex Fluorescent Immunohistochemistry (mfIHC)**

Multiplex fluorescent IHC was carried out on a Ventana Discovery Ultra (Roche) using tyramide signal amplification (TSA; Akoya Biosciences, Marlborough, MA, USA). Each target was sequentially labeled using horseradish peroxidase (HRP)-conjugated secondary antibodies and corresponding Opal fluorophores, followed by heat-induced stripping between rounds to remove antibody complexes while preserving fluorescent signal.For all staining panels, following primary antibody incubation, slides were treated with a goat anti-rabbit HRP-polymer secondary antibody (Vector Laboratories, Burlingame, CA, Cat# VP1102) for 20 minutes at 37 °C prior to fluorophore development. This step was repeated for each round of staining.

***Acute lung tissue panel:*** Following CC1 pretreatment (Roche Cat# 950-500, 95 °C for 32 minutes), slides were incubated with Rabbit anti-SARS-CoV-2 S1 (1:400, CST Cat# 99423), Rabbit anti-CD11b (1:2000, Abcam Cat# ab13357), Rabbit anti-CD8 (1:100, CST Cat# 98941), Rabbit anti-CD3e (1:100, Dako Cat# A045201-2), Rabbit anti-Iba1 (1:250, CST Cat# 17198). Primary antibodies were incubated at 37 °C for 32–60 minutes, followed by HRP-conjugated goat anti-rabbit or anti-mouse secondary antibodies (Vector Laboratories) and Opal fluorophore development (20 min per fluor). Opals used: Opal 570 (1:250, for S1), Opal 520 (1:120, for CD11b), Opal 620 (1:60, for CD8), Opal 480 (1:100, for CD3e), Opal 690 (1:125, for Iba1). All slides were counterstained with spectral DAPI (Akoya Biosciences) and mounted with ProLong Gold Antifade Reagent (Thermo Fisher Scientific).

***Post-acute lung (30 & 65 dpi):*** For post-acute lung, CC1 (95 °C for 1 hour) antigen retrieval was performed prior to antibody staining. Samples were then stained with the following antibodies: Rabbit anti-CD19 (1:400, CST Cat# 90176), Rabbit anti-CD11b (1:2000, Abcam Cat# ab13357), Rabbit anti-CD8 (1:100, CST Cat# 98941), Rabbit anti-CD4 (1:150, CST Cat# 25229), Rabbit anti-Iba1 (1:250, CST Cat# 17198). Primary antibodies were incubated at 37 °C for 32–60 minutes. For detection, the following fluorophores were used: Opal 570 (1:90, CD8), Opal 520 (1:80, CD19), Opal 620 (1:90, CD11b), Opal 480 (1:100, CD4), Opal 690 (1:115, Iba1). All slides were counterstained with spectral DAPI (Akoya Biosciences) and mounted with ProLong Gold Antifade Reagent (Thermo Fisher Scientific).

***Post-acute brain (30 & 65 dpi):*** For post-acute brain, samples were stained with the following antibodies: Rabbit anti-GFAP (1:100, CST Cat# 80788), Rabbit anti-NeuN (1:150, CST Cat# 24307), Rabbit anti-CD8 (1:100, CST Cat# 98941), Rabbit anti-CD3e (1:100, Invitrogen Cat# MA5-14524), Rabbit anti-CD11b (1:2000, Abcam Cat# ab13357), Rabbit anti-Iba1 (1:250, CST Cat# 17198). For detection the following fluorophores were used: Opal 570 (1:100, CD8), Opal 520 (1:100, NeuN), Opal 620 (1:225, GFAP), Opal 480 (1:120, CD11b), Opal 690 (1:90, CD3e), Opal 780 (1:35, Iba1). All slides were counterstained with spectral DAPI (Akoya Biosciences) and mounted with ProLong Gold Antifade Reagent (Thermo Fisher Scientific).

**RNA In Situ Hybridization (ISH) for Human ACE2**

RNAscope ISH was performed using the RNAscope 2.5 LSx Reagent Kit (Advanced Cell Diagnostics, Newark, CA, USA) on the BOND RXm platform (Leica Biosystems). Sections were hybridized with an anti-sense probe specific to human ACE2 (NM_021804.3; Cat# 848038), which lacks cross-reactivity to murine Ace2. Amplification steps included incubation with AMP 5 and AMP 6 buffers for 45 and 30 minutes, respectively. Vero E6 cells were used as positive controls, and murine Ppib mRNA served as a quality control.

**Whole-Slide Imaging and Spectral Unmixing**

Fluorescently stained slides were imaged on a PhenoImager HT (Akoya Biosciences). Exposure settings for each Opal fluorophore were calibrated based on regions of high signal intensity to minimize noise. Spectral unmixing and autofluorescence removal were performed using InForm software (v3.0, Akoya), applying dye-specific spectral libraries.

**Image Analysis**

All image analysis was conducted in HALO (Indica Labs, Corrales, NM) using the following modules:

***Acute Lung:*** Quantification of SARS-CoV-2 S immunoreactivity and Iba1 area was performed using the Area Quantification (AQ) module (v2.1.11). Regions of interest were annotated, and signal thresholds were adjusted in real time to capture stain-specific intensity. Lung consolidation was quantified using a Random Forest tissue classifier trained on manually annotated examples of consolidated and non-consolidated regions. Quantification of CD11b+, CD8+, and CD3e+ cells was performed using the HighPlex FL module (v4.0.4). Cells were segmented based on DAPI-stained nuclei, with nuclear size and shape thresholds adjusted to match ground truth segmentation. Marker positivity was defined using minimum fluorophore intensity thresholds for cytoplasmic compartments, customized per marker and sample using real-time tuning. Output data included absolute cell counts, percentages per phenotype, and area analyzed, exported as .CSV files.

***Post-Acute Lung:*** HP module parameters were reused from the acute lung analysis and adapted as needed. Cell types quantified included CD19+, CD11b+, CD8+, CD4+, and CD3e+ populations. Output data included absolute cell counts, percentages per phenotype, and area analyzed, exported as .CSV files.

***Post-Acute Brain:*** Tissue regions were annotated using HALO's flood-fill tool. The AQ module (v2.3.4) quantified signal area for CD11b, Iba1, GFAP, and dual CD11b+Iba1+ expression. Cellular quantification of NeuN+, CD8+, and CD3e+ populations was performed using HighPlex FL (v4.2.3) as above. Marker positivity was defined using minimum fluorophore intensity thresholds for both nuclear and cytoplasmic compartments, customized per marker and sample using real-time tuning. Output data included absolute cell counts, percentages per phenotype, and area analyzed, exported as .CSV files.

#### SUPPLEMENTAL FIGURES AND LEGENDS

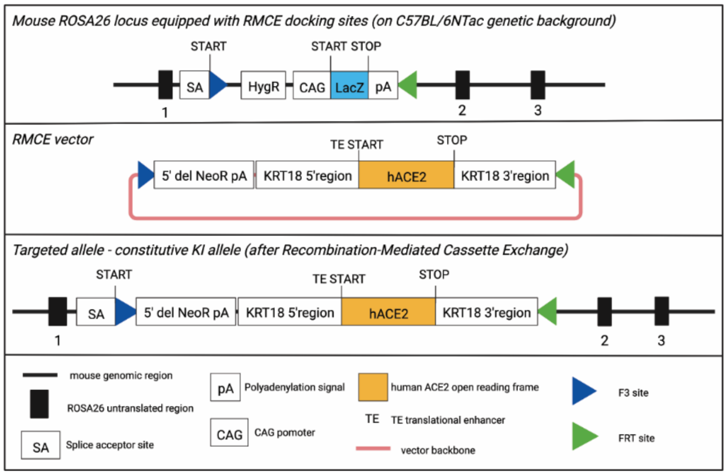


**Figure S1. Schematic showing the strategy used to generate the targeted transgenic ACE2 knock-in allele in RAB/6N mice.**

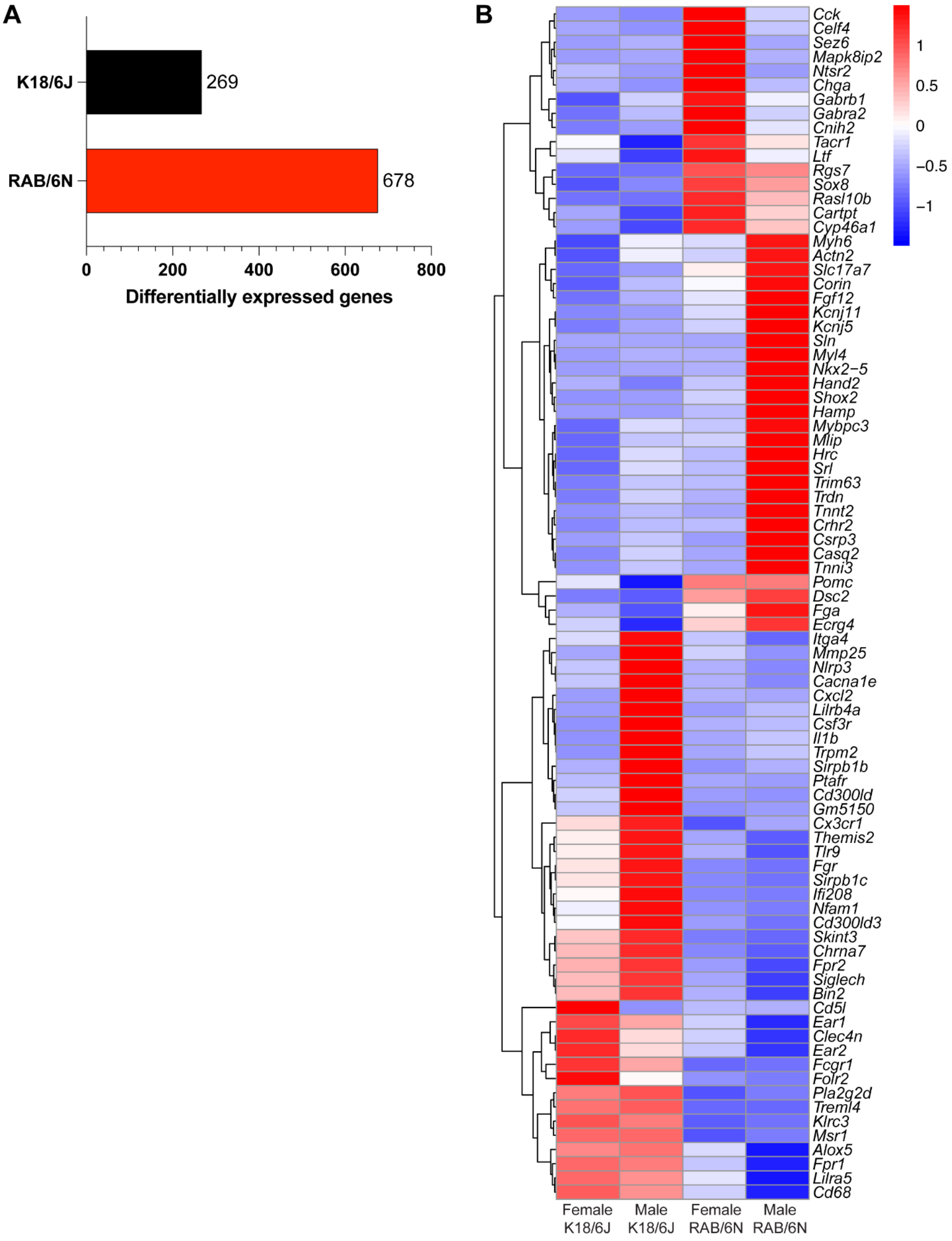
A
K18/6J
269
RAB/6N
678
0
200
400
600
800
Differentially expressed genes
B
Cck
Celf4
Sez6
Mapk8ip2
Ntsr2
Chga
Gabrb1
Gabra2
Cnih2
Tacr1
Ltf
Rgs7
Sox8
Rasl10b
Cartpt
Cyp46a1
Myh6
Actn2
Slc17a7
Corin
Fgf12
Kcnj11
Kcnj5
Sln
Myl4
Nkx2−5
Hand2
Shox2
Hamp
Mybpc3
Mlip
Hrc
Srl
Trim63
Trdn
Tnnt2
Crhr2
Csrp3
Casq2
Tnni3
Pomc
Dsc2
Fga
Ecrg4
Itga4
Mmp25
Nlrp3
Cacna1e
Cxcl2
Lilrb4a
Csf3r
Il1b
Trpm2
Sirpb1b
Ptafr
Cd300ld
Gm5150
Cx3cr1
Themis2
Tlr9
Fgr
Sirpb1c
Ifi208
Nfam1
Cd300ld3
Skint3
Chrna7
Fpr2
Siglech
Bin2
Cd5l
Ear1
Clec4n
Ear2
Fcgr1
Folr2
Pla2g2d
Treml4
Klrc3
Msr1
Alox5
Fpr1
Lilra5
Cd68
1
0.5
0
−0.5
−1
Female K18/6J
Male K18/6J
Female RAB/6N
Male RAB/6N

**Figure S2. K18-hACE2/6J and RAB/6N show distinct lung transcriptomes at baseline. (A)** Number of genes upregulated in K18-hACE2/6J and RAB/6N lungs. **(B)** Heatmap showing select, significantly upregulated genes in lungs of male and female naïve RAB/6N and K18-hACE2/6J mice. male n=3, female n =4.

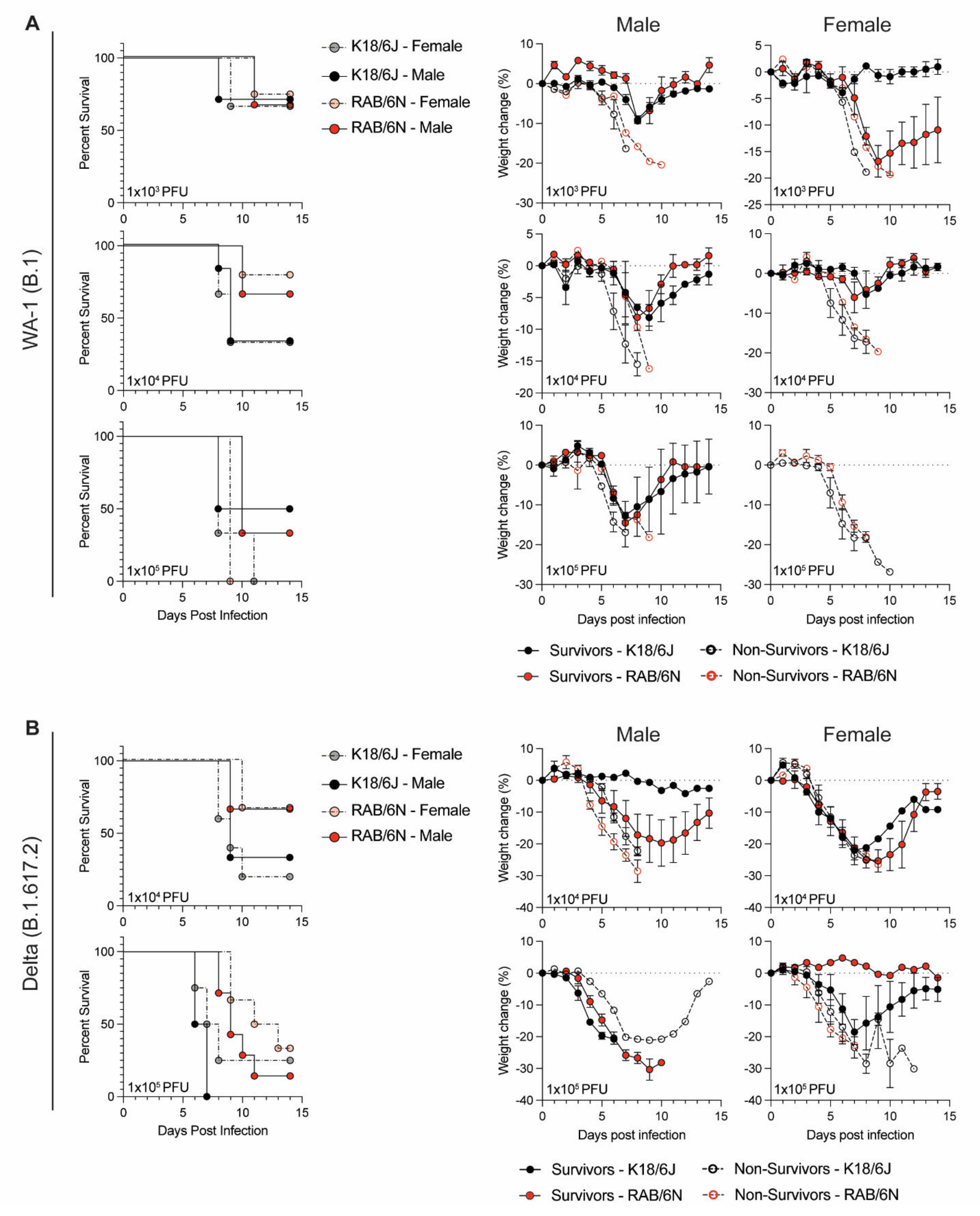
A
WA-1 (B.1)
K18/6J - Female
K18/6J - Male
RAB/6N - Female
RAB/6N - Male
Male
Female
Percent Survival
Weight change (%)
1x10³ PFU
1x10⁴ PFU
1x10⁵ PFU
Days Post Infection
Days post infection
Survivors - K18/6J
Survivors - RAB/6N
Non-Survivors - K18/6J
Non-Survivors - RAB/6N
B
Delta (B.1.617.2)

**Figure S3. Male and female weight loss dynamics after SARS-CoV-2 challenge.** 12–22-week-old male and female RAB/6N (red) and K18-hACE2/6J mice (black) were infected intranasally with SARS-CoV-2 WA-1 **(A)** and Delta **(B)** and monitored for 14 days. Weight loss and survival data was analyzed for males and females independently. For weight loss, open circles indicate mice that succumb to infection and filled circles represent mice that survived infection. For survival, semi-transparent circles represent females and non-transparent represent male. RAB/6N mice are in red and K18-hACE2/6J mice are in black.

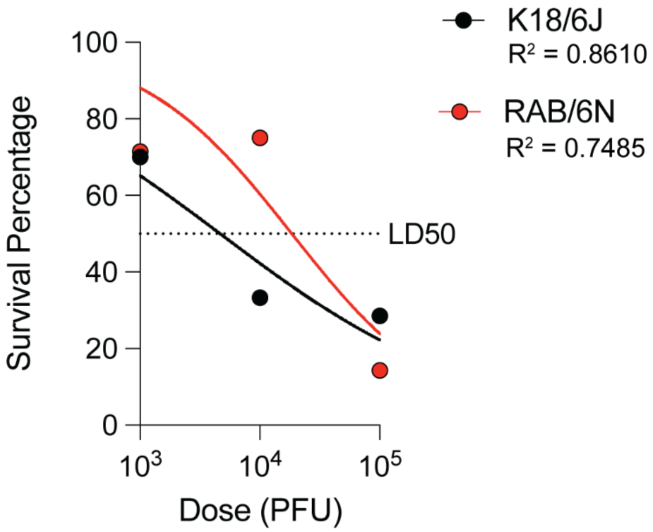


**Figure S4. Correlation plot of dose and percent lethality in RAB/6N and K18-hACE2/6J mice.** Points represent survival rate at each viral dose. Lines indicate the non-linear regression curve. Dotted line represents the viral dose to achieve 50% lethality (LD50).

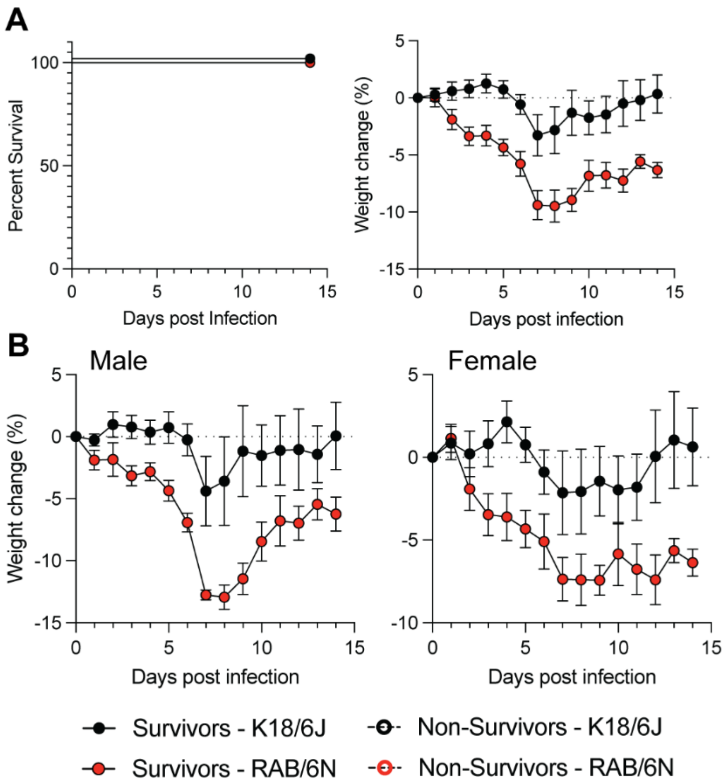


**Figure S5. Omicron BA.1 infection causes sex-biased disease.** 12–22-week-old male and female RAB/6N (red) and K18-hACE2/6J mice (black) were infected intranasally with $1x10^4$ pfu of SARS-CoV-2 Omicron BA.1. **(A)** Survival plot of RAB/6N and K18-hACE2/6J (left) and average weight loss (right). A Mantel-Cox analysis was performed to assess significance of survival. **(B)** Weight loss of male (left) and female (right) RAB/6N (red) and K18-hACE2/6J (black) mice.

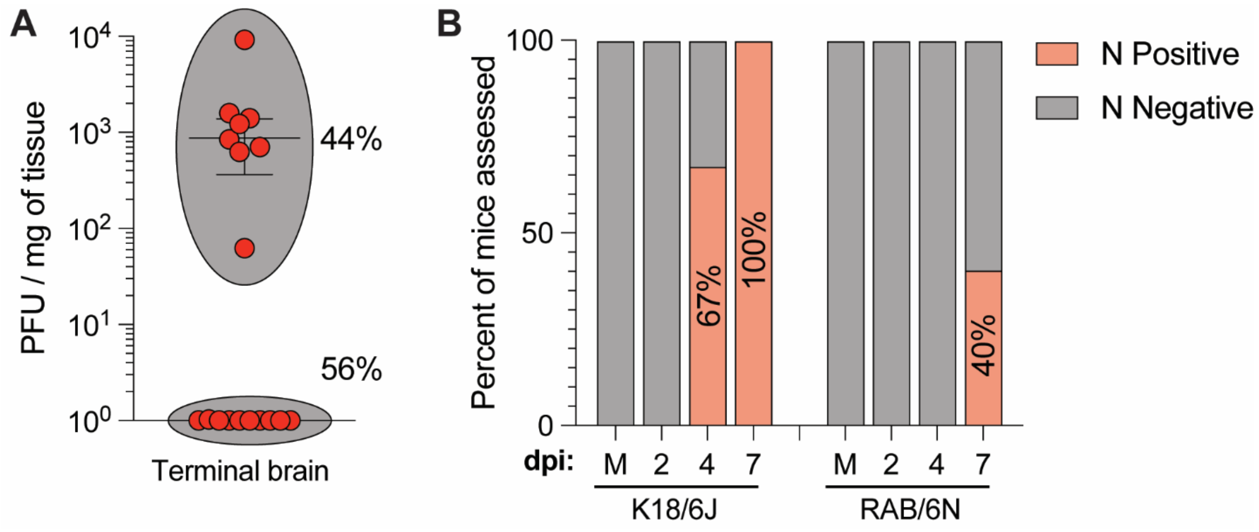


**Figure S6. Quantification of neuroinvasion in RAB/6N mice. (A)** RAB/6N mice were infected with 1x10$^5$ pfu of SARS-CoV-2 Delta. At euthanasia criteria, brains were collected and viral titers were quantified by plaque-forming assay. Weight is displayed as pfu/mg of brain tissue. **(B)** RAB/6N mice were infected with 1x10$^5$ pfu of SARS-CoV-2 WA-1 and euthanized at 2, 4, or 7 dpi, along with non-infected (mock, M) mice. IHC for SARS-CoV-2 N was then performed on brain tissue section to establish the percentage of mice positive for viral antigen in the brain. Bar chart indicates the percentage of N antigen positive brains at each time points for K18-hACE2/6J and RAB/6N mice.

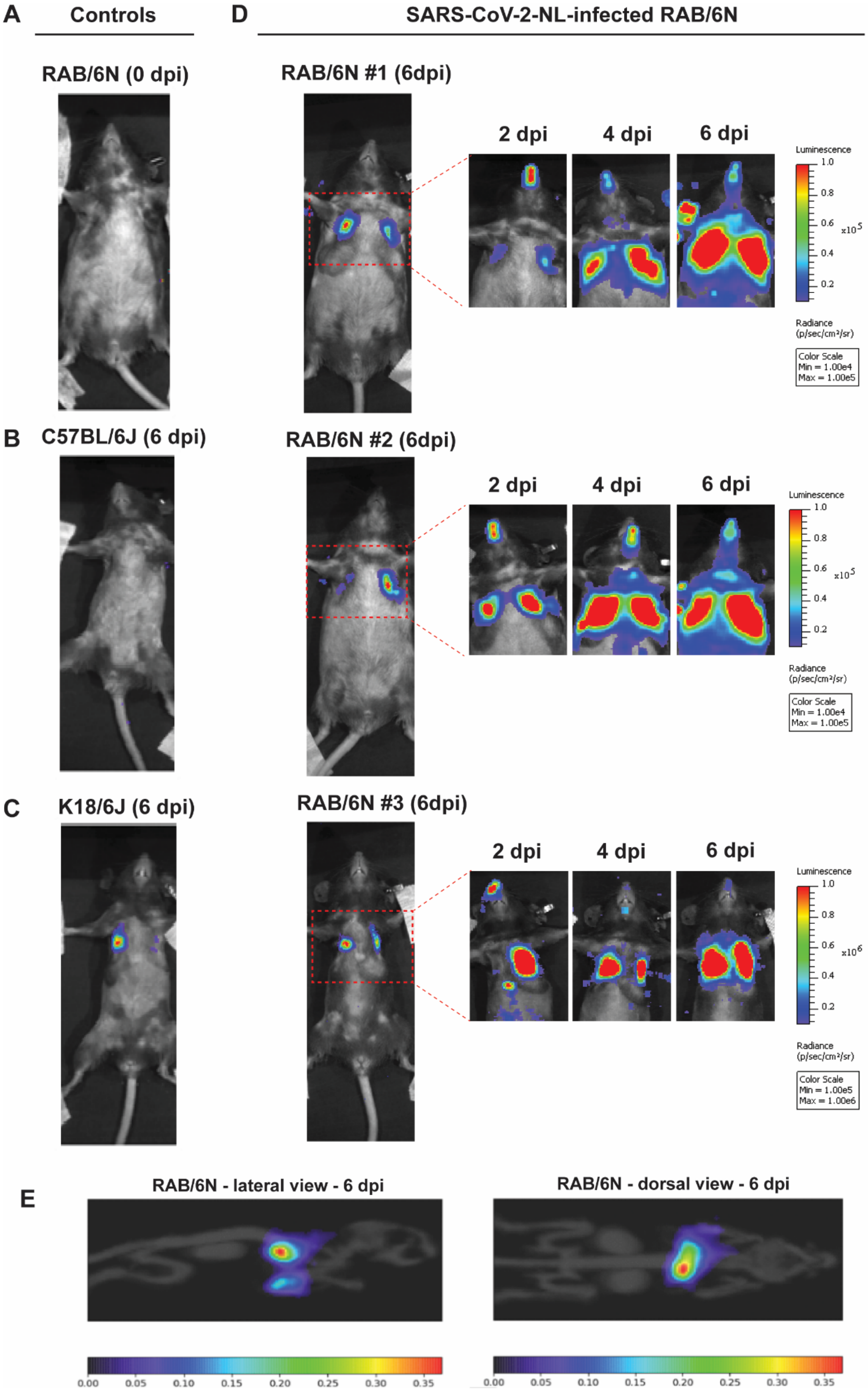
A
Controls
RAB/6N (0 dpi)
B
C57BL/6J (6 dpi)
C
K18/6J (6 dpi)
D
SARS-CoV-2-NL-infected RAB/6N
RAB/6N #1 (6dpi)
2 dpi
4 dpi
6 dpi
Luminescence
Radiance (p/sec/cm²/sr)
Color Scale Min = 1.00e4 Max = 1.00e5
RAB/6N #2 (6dpi)
RAB/6N #3 (6dpi)
Color Scale Min = 1.00e5 Max = 1.00e6
E
RAB/6N - lateral view - 6 dpi
RAB/6N - dorsal view - 6 dpi

615

**Figure S7. *In vivo* imaging of RAB/6N mice reveals sustained infection.** K18-hACE2/6J, RAB/6N, and C57BL/6J mice were infected with $1x10^5$ PFU of SARS-CoV-2 harboring NanoLuciferase and imaged using a PerkinElmer IVIS Spectrum at 0, 2, 4, and 6 dpi. **(A-C)** Images of a control RAB/6N mouse at 0 dpi (prior viral challenge, A), C57BL/6J infected mouse at 6 dpi (B), and K18-hACE2/6J mouse at 6 dpi (C) are shown. **(D)** Images of three RAB/6N mice acquired at 2, 4, and 6 dpi. Whole body images were taken at 6 dpi. **(E)** lateral and dorsal view of a representative RAB6/N mice imaged at 6 dpi. Images were acquired using the InVivo Analytics InvivoPLOT™ mirror gantry.

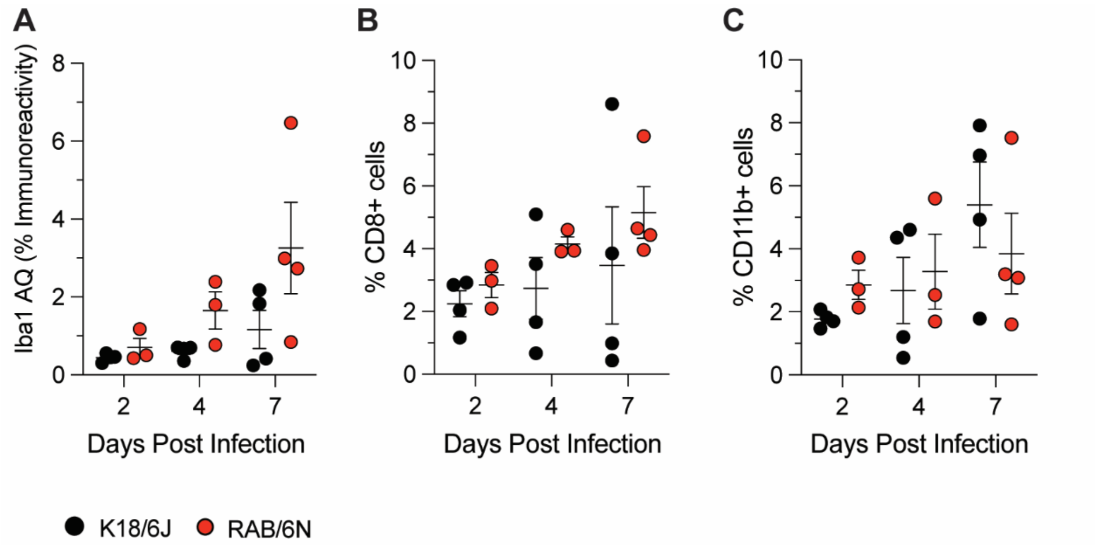


**Figure S8. Halo quantification of immune populations from multiplex immunohistochemistry.** Halo analysis software was used to quantify diverse hematopoietic populations from whole slide images of lungs from RAB/6N and K18-hACE2/6J mice at 2, 4 and 7 dpi. **(A)** Area quantification (AQ) of Iba1+ positive cells. **(B-C)** Percent of CD8+ (B) and CD11b+ positive cells (C) among total cells in analyzed lung tissue sections following mIHC. Dot plots indicate individual mice. Error bars indicated mean ± standard error of the mean (SEM).

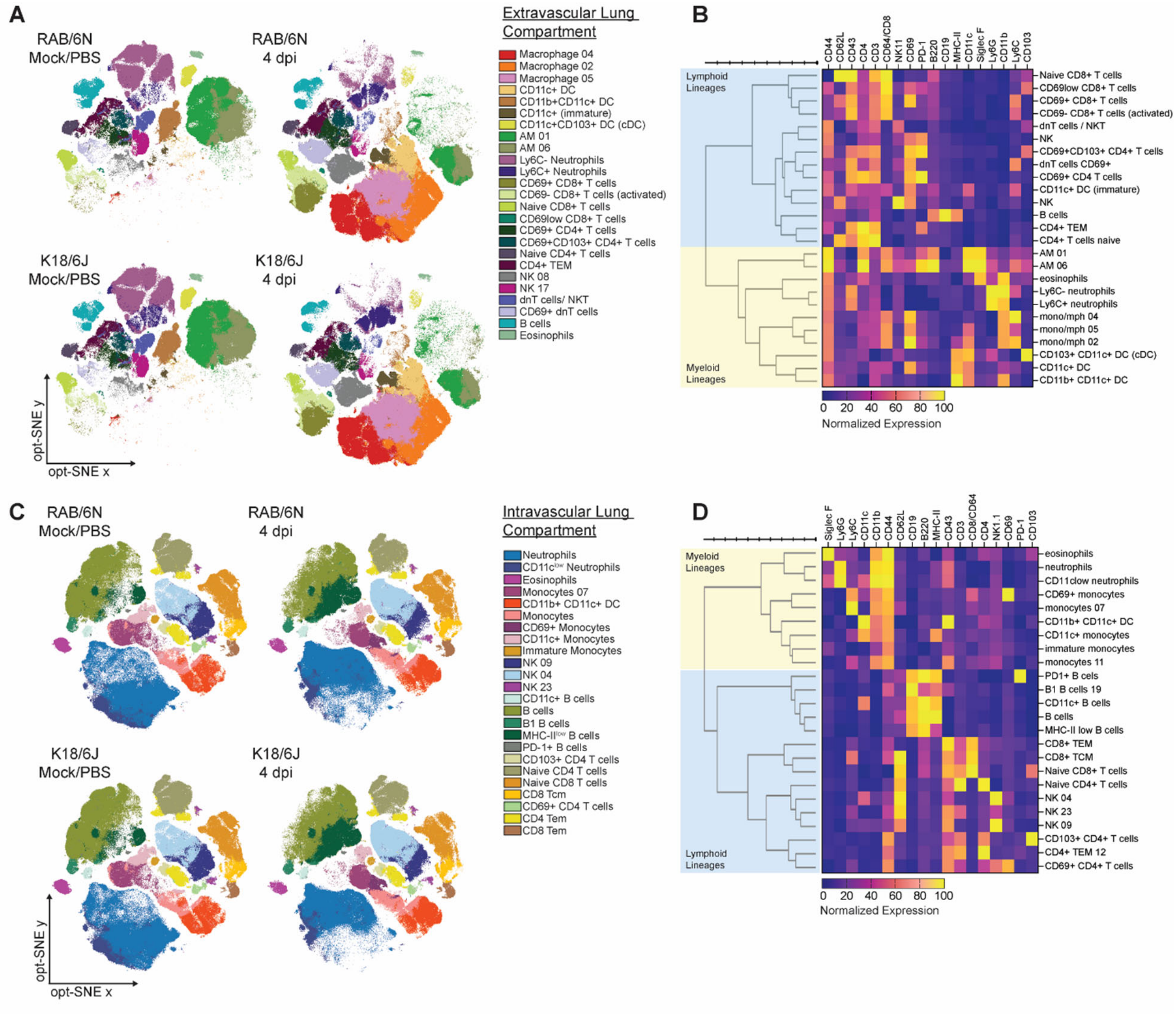


**Figure S9. Unbiased immune profiling of the lung immune environment by flow cytometry.** K18-hACE2/6J and RAB/6N mice were infected with 1x10$^{5}$ pfu of SARS-CoV-2 WA-1 isolate or mock infected, and immune populations in the lung were assessed at 4 dpi. **(A)** Lung extravascular (i.v. CD45.2$^{neg}$CD45$^{pos}$) immune compartment in mock (left) and 4 dpi (right) spectral cytometry data projected into opt-SNE space and clustered with Phenograph. **(B)** Heatmap showing normalized expression of each marker in Phenograph clusters. Myeloid lineages are marked by yellow and lymphocyte lineages are marked by blue. Clusters were annotated based on phenotype. **(C)** Lung intravascular (i.v. CD45.2$^{pos}$CD45$^{pos}$) immune compartment in mock (left) and 4 dpi (right) projected into opt-SNE space and clustered with Phenograph. **(D)** Heatmap showing normalized expression of each marker in Phenograph clusters. Myeloid lineages are marked by yellow and lymphocyte lineages are marked by blue. Clusters were annotated based on phenotype.

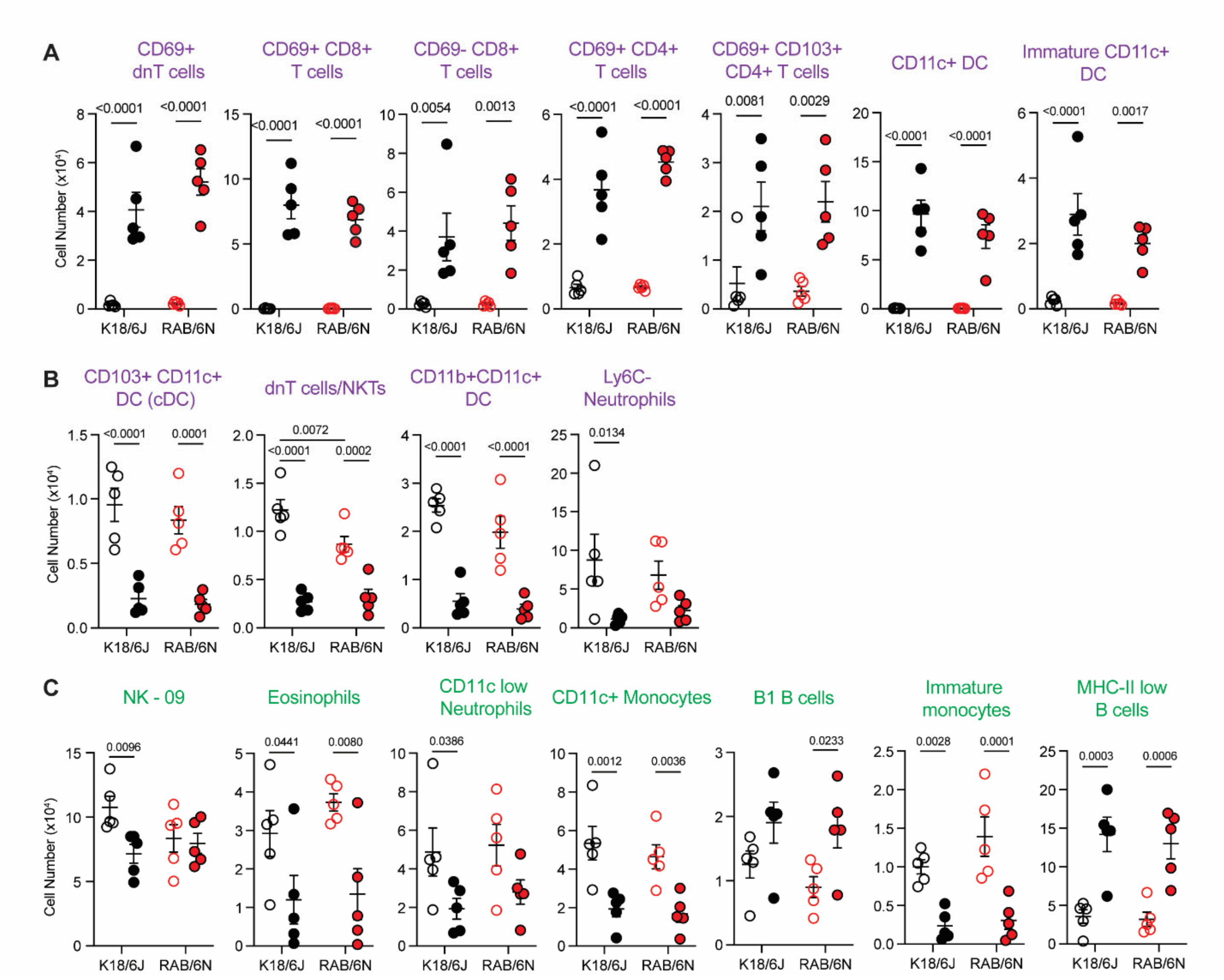


**Figure S10. Immune populations in the lung that trend similarly in K18-hAEC2/6J and RAB/6N mice. (A-B)** Lung intravascular immune populations that **(A)** increased or **(B)** decreased in cell numbers in both K18-hACE2/6J and RAB/6N mice after infection. **(C)** Lung extravascular immune populations that increased or decreased in both K18-hACE2/6J and RAB/6N mice after infection.

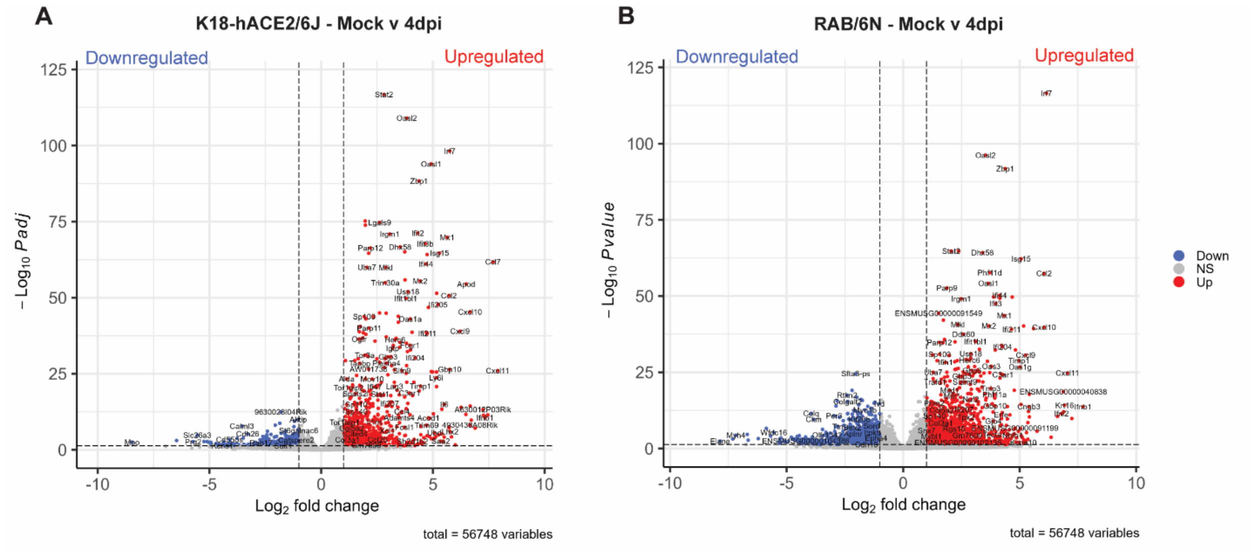


**Figure S11. K18-hACE2/6J and RAB/6N mice show robust transcriptional responses in the lung at 4 days post infection.** Volcano plots highlighting upregulated and downregulated genes in the lungs of K18-hACE2/6J **(A)** and RAB/6N **(B)** mice between non-infected (mock) mice and 4 dpi. Significantly upregulated genes are in red and significantly downregulated genes are in blue. Non-significant genes are in grey.

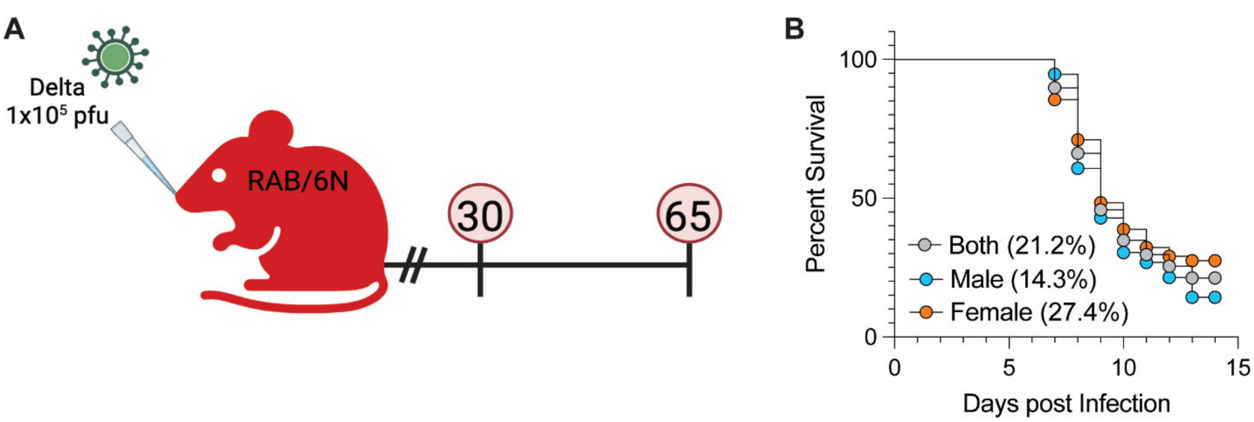


**Figure S12. Development of PASC model from mice surviving infection with Delta. (A)** RAB/6N mice were infected with $1x10^5$ pfu of SARS-CoV-2 Delta and mice that survived acute disease were euthanized at 30 or 65 dpi to characterize long-term disease. Created with BioRender.com. **(B)** Survival plot of Delta-infected mice showing the average survival of all mice (21.2%; grey), female (27.4%; orange), and male mice (14.3%; blue).

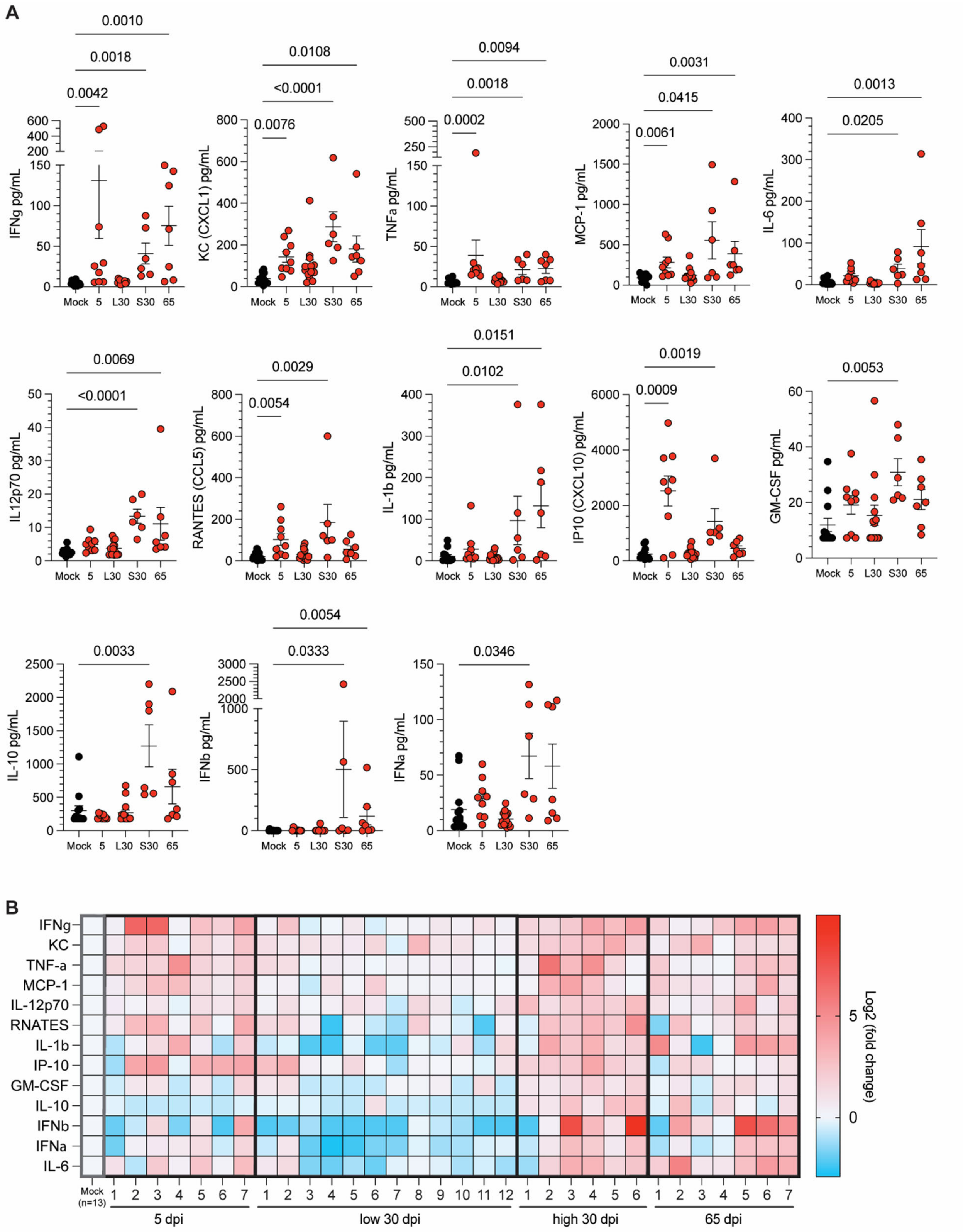
A
IFNg pg/mL
0.0010
0.0018
0.0042
KC (CXCL1) pg/mL
0.0108
<0.0001
0.0076
TNFa pg/mL
0.0094
0.0018
0.0002
MCP-1 pg/mL
0.0031
0.0415
0.0061
IL-6 pg/mL
0.0013
0.0205
IL12p70 pg/mL
0.0069
<0.0001
RANTES (CCL5) pg/mL
0.0029
0.0054
IL-1b pg/mL
0.0151
0.0102
IP10 (CXCL10) pg/mL
0.0019
0.0009
GM-CSF pg/mL
0.0053
IL-10 pg/mL
0.0033
IFNb pg/mL
0.0054
0.0333
IFNa pg/mL
0.0346
Mock
5
L30
S30
65
B
IFNg
KC
TNF-a
MCP-1
IL-12p70
RNATES
IL-1b
IP-10
GM-CSF
IL-10
IFNb
IFNa
IL-6
Mock (n=13)
5 dpi
low 30 dpi
high 30 dpi
65 dpi
Log2 (fold change)

**Figure S13. RAB/6N$^{PASC}$ mice show persistent systemic inflammation. (A)** Dot plots indicating individual cytokine levels in non-infected (mock) mice and infected mice at 5, 30 (severe: s and low: l inflammation), and 65 dpi. Error bars indicated mean ± standard error of the mean (SEM). n=7-9 **(B)** Heatmap showing $\log_2$ fold change of cytokine levels in individual acutely infected and RAB/6N$^{PASC}$ mice compared to non-infected (mock) mice. Each column represents an individually infected mice at the exception of the mock column, which represents an average of 13 non-infected (mock) mice.

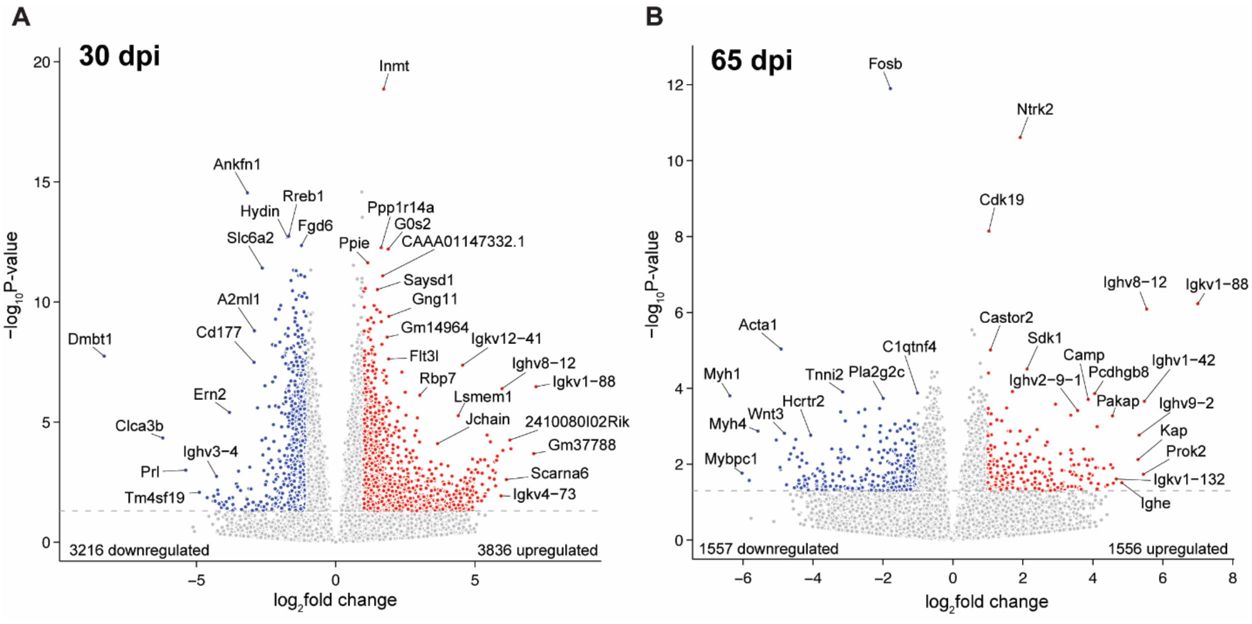


**Figure S14. RAB/6N PASC mice show signs of multi-tissue gene dysregulation. (A-B)** Volcano plots highlighting upregulated and downregulated genes in the lung of RAB/6N mice at **(A)** 30 dpi and **(B)** 65 dpi. Significantly upregulated genes are in red and significantly downregulated genes are in blue. Non-significant genes are in grey.

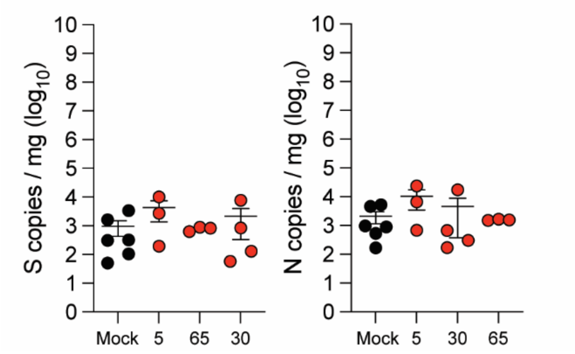


**Figure S15. RAB/6N$^{PASC}$ mice are free of viral RNA in heart tissues.** RT-qPCR analysis of SARS-CoV-2 S and N in the heart of RAB$^{PASC}$ mice during acute infection (5 dpi) and post-acute infection (30 and 65 dpi) and in non-infected (mock) mice. Dot plots with error bars indicating mean ± standard error of the mean (SEM).

| Tissue | DPI | Gene Name | Regu-lation | Associated effects | Associated Diseases | References |
|---|---|---|---|---|---|---|
| **LUNG - 30DPI** | | | | | | |
| Lung | 30 | *Igkv/hv/ Jchain* | Up | Increased Ig production | Inflammation, autoimmunity | *(33-37)* |
| Lung | 30 | *Flt3L* | Up | Stromal and dendritic cell cytokine support | Chronic inflammation, autoimmunity, predisposition to fibrosis | (*38*) |
| Lung | 30 | *G02s* | Up | T-cell proliferation | | (*39*) |
| Lung | 30 | *Gng11* | Up | Cellular senescence | Predisposition to fibrosis, impaired susceptibility to infections, potential oncogenic shifts | (*40*) |
| Lung | 30 | *CD177* | Down | Impaired neutrophil activation and migration | Increased susceptibility to infection, immune tolerance | (*41*) |
| Lung | 30 | *DMBT1* | Down | Dysregulation of epithelial barrier homeostasis | Allergic airway diseases, interstitial lung diseases, COPD-like diseases | (*42*) |
| Lung | 30 | *Clca3b* | Down | | | *(43)* |
| Lung | 30 | *Hydin* | Down | Defective cilia motility and mucociliary clearance | COPD-like diseases, bronchiectasis | (*44*) |
| Lung | 30 | *Ern2* | Down | Decreased goblet cell development and mucus barrier assembly | Increased susceptibility to infection, bronchiectasis , mucus differentiation disorders | (*45, 46*) |
| **LUNG - 65DPI** | | | | | | |

| | | | | | | |
|---|---|---|---|---|---|---|
| Lung | 65 | *Igkv/hv/ Jchain* | Up | Increased Ig production | Chronic inflammation, autoimmunity | *(33-36)* |
| Lung | 65 | *IgE* | Up | Allergy/Type2 autoimmunity | | (*47-51*) |
| Lung | 65 | *CAMP* | Up | Upregulation of antimicrobial responses | | (*52*) |
| Lung | 65 | *Castor2* | Up | Negative regulation of mTORC1 pathway, metabolic stress | Metabolic and immune disorders, predisposition to fibrosis | (*53*) |
| Lung | 65 | *KAP* | Up | Increased oxidative stress | Fibrosis, hypertension | (*54*) |
| Lung | 65 | *CDK19* | Up | Hypertrophic growth | Cancers | (*55*) |
| Lung | 65 | *Ntrk2* | Up | Enhanced lung-CNS crosstalks, cellular survival and differentiation | Lung cancer | *(56)* |
| Lung | 65 | *Prok2* | Up | Enhanced lung-CNS crosstalks, macrophage chemotaxis and phagocytosis, regulation of circadian clock, metabolism and olfaction (anosmia) | Chronic inflammation, olfactory defects, metabolic disorders | *(57-61)* |
| Lung | 65 | *Sdk1* | Up | Enhanced cell adhesion, proliferation | Cancers, Lung adenocarcino ma | *(62)* |
| Lung | 65 | *Pcdhgb 8* | Up | | | (*63, 64*) |
| Lung | 65 | *Fobs* | Down | Downregulation of IL-17 responses | Increased susceptibility to infections | (*65-69*) |
| Lung | 65 | *Hctr2* | Down | | | (*70*) |
| Lung | 65 | *C1qTN F4* | Down | Decreased activation of the STAT3 and NF-κB pathways and cell survival | Increased susceptibility to infections, predisposition to SLE | *(71, 72)* |

| | | | | | | |
|---|---|---|---|---|---|---|
| Lung | 65 | *Pla2g2c* | Down | Dysregulation of inflammation via decreased immune mediator activation, impaired clearance of immune complexes | Increased susceptibility to infections, autoimmune disorders | *(73, 74)* |
| Lung | 65 | *Wnt3* | Down | Defective tissue repair | Cancers | (*75*) |
| Lung | 65 | *Acta1* | Down | Impaired cytoskeletal remodeling, loss of resident stromal and mesenchymal cells | | (*76*) |
| Lung | 65 | *Myh1/4* | Down | | | (*77*) |
| Lung | 65 | *Mybpc1* | Down | | | (*78*) |
| **BRAIN - 30DPI** | | | | | | |
| Brain | 30 | *Igkv/hg/hv/kc/lv* | Up | Increased Ig production | Inflammation, auto-inflammation, brain and cognitive damage | *(33-36)* |
| Brain | 30 | *Inmt* | Up | Enhanced DMT production, response to cerebral ischemia | Protective mechanisms in response to inflammation and cellular/tissue stress? | (*79-81*) |
| Brain | 30 | *Neurod6* | Up | Neuronal protection, differentiation and survival | | (*82, 83*) |
| Brain | 30 | *Sim1* | Down | Hypothalamic neuroendocrine dysfunction | Mood disorders, depression, obesity | *(84)* |
| Brain | 30 | *Cartpt* | Down | | | *(85)* |
| Brain | 30 | *Ucn* | Down | | | (*86*) |
| Brain | 30 | *Prg2* | Down | Defective eosinophil degranulation | Cancer, brain infection, neurological deficits | (*87-92*) |
| Brain | 30 | *Epx* | Down | | | |
| Brain | 30 | *Ear6* | Down | Defective eosinophil ribonucleolytic functions | | (*89, 90, 93*) |
| Brain | 30 | *Mfsd2a* | Down | Blood-brain barrier defects | Brain infection, cognitive impairment, neuro-inflammation, neuro- | (*94-99*) |
| Brain | 30 | *Slco1b2* | Down | Impaired neuronal transport of molecules/hormones | | (*100-102*) |

| | | | | | | |
|---|---|---|---|---|---|---|
| | | | | | degenerative disorders | |
| **HEART - 65DPI** | | | | | | |
| Heart | 65 | *Gp9* | Up | Megakaryocyte and platelet infiltration | Chronic inflammation, Myocarditis, thrombo-inflammatory cardio-myopathy | *(103)* |
| Heart | 65 | *Itga2* | Up | | | |
| Heart | 65 | *Tubb1* | Up | | | *(104)* |
| Heart | 65 | *Clec1b* | Up | | | *(105)* |
| Heart | 65 | *Treml1* | Up | | | *(106)* |
| Heart | 65 | *Igkv8-24* | Up | Ig production | | *(33-36)* |
| Heart | 65 | *cd266* | Up | T-cell activation | | *(107)* |
| Heart | 65 | *Ctla2a* | Up | T-cell exhaustion | | *(108)* |
| Heart | 65 | *Cysltr2* | Up | Type 2 immunopathology | | *(109)* |
| Heart | 65 | *Phgdh* | Up | Metabolic stress | Cancer | *(110)* |
| Heart | 65 | *Saa3* | Up | | Obesity, diabetes, and cardio-vascular disease | *(111)* |
| Heart | 65 | *Col3a1* | Up | Collagen I fibrillogenesis | Heart Fibrosis | *(112)* |
| Heart | 65 | *Bnip3* | Up | Increased mitochondrial membrane permeabilization and cell death | Heart stress and failure | *(113, 114)* |
| Heart | 65 | *Tspo* | Up | Impaired mitochondria energy production, Ca2+ homeostasis, and mitophagy. Increased ROS production and inflammation | Associated with cancers, Alzheimer and Parkinson disease, inflammation-induced cardiac dysfunctions. | *(114-117)* |
| Heart | 65 | *Dbi* | Up | Altered lipid metabolism, enhanced cell survival, low-grade inflammation | Chronic inflammation, cancers, increased susceptibility to infections | *(118)* |
| Heart | 65 | *Heyl* | Down | Impaired angiogenesis | Advanced heart failure, ischemic | *(119)* |
| Heart | 65 | *Fgf6* | Down | | | *(120)* |
| Heart | 65 | *Srf* | Down | | | *(121)* |

| | | | | | | |
|---|---|---|---|---|---|---|
| Heart | 65 | *Apln* | Down | Impaired cardiomyocyte differentation and contractility | cardio-myopathy | *(122)* |
| Heart | 65 | *Bcl6b* | Down | Decreased cell survival, lymphocyte activation | Increased tissue susceptibility to cardiac stress, heart damage and failure | (*123, 124*) |
| Heart | 65 | *Hspb6* | Down | Decreased protection from oxydative stress | | (*125*) |
| Heart | 65 | *Cyp8b1* | Down | Impaired lipid signaling and liver-heart crosstalk | Cardio-metabolic diseases, heart failure | *(126)* |
| Heart | 65 | *S1pr3* | Down | Blunted sphingolipid signaling, myeloid differentiation, vascular repair and tone, cardiomyocyte integrity | | *(127-129)* |
| Heart | 65 | *Tnfa* | Down | Reduced protection against cardiac myocarditis and cardiac death | Post-inflammatory cardio-myopathy, heart failure | (*130-132*) |

**Supplemental Table 1. List of select genes differentially regulated in RAB/6N$^{PASC}$ mice with associated literature references.** Select genes significantly up- (light red) or downregulated (light blue) in the lung, brain and heart of RAB/6N mice at 30 and/or 65 dpi. Associated effects and disease phenotypes are listed for each gene showing statistically significant differential regulation, along with the supporting literature references.
